# A new monoclonal antibody enables BAR analysis of subcellular importin β1 interactomes

**DOI:** 10.1101/2022.03.23.485495

**Authors:** Didi-Andreas Song, Stefanie Alber, Ella Doron-Mandel, Vera Schmid, Christin A. Albus, Orith Leitner, Hedva Hamawi, Juan A. Oses-Prieto, Alma L. Burlingame, Mike Fainzilber, Ida Rishal

## Abstract

Importin β1 (KPNB1) is a nucleocytoplasmic transport factor with critical roles in both cytoplasmic and nucleocytoplasmic transport, hence there is keen interest in the characterization of its subcellular interactomes. We found limited efficiency of BioID in detection of importin complex cargos, and therefore generated a highly specific and sensitive anti-KPNB1 monoclonal antibody to enable Biotinylation by Antibody Recognition (BAR) analysis of importin β1 interactomes. The monoclonal antibody recognizes an epitope comprising residues 301-320 of human KPBN1, and strikingly is highly specific for cytoplasmic KPNB1 in diverse applications, with little or no reaction with KPNB1 in the nucleus. BAR with this novel antibody revealed numerous new interactors of importin β1, expanding the KPNB1 interactome to cytoplasmic and signaling complexes that highlight potential new functions for the importins complex beyond nucleocytoplasmic transport. Data are available via ProteomeXchange with identifier PXD032728.

## Introduction

Transport across the nuclear envelope is an active receptor-mediated process, which is essential for normal cell function (1). The importin β protein family transports cargos with nuclear localization signals (NLS) into the nucleus. Nearly all other importin βs bind their cargos directly, while the canonical importin β1 (also termed karyopherin β1, KPNB1) uses one of the accessory importin α family members as cargo binding subunits (2). Ran-regulated association of importin β1/α/NLS-cargo complexes in the cytoplasm and their dissociation in the nucleoplasm enables nucleocytoplasmic transport (Figure 1A). In addition to its key role in nucleocytoplasmic transport, importin β1 forms cytoplasmic transport complexes with the microtubule motor dynein (Figure 1B). Importin β1 dependent cytoplasmic transport has been implicated in long distance signaling in large cells, especially neurons, with roles in injury signaling, size regulation and growth control (3, 4).

**Figure 1:**
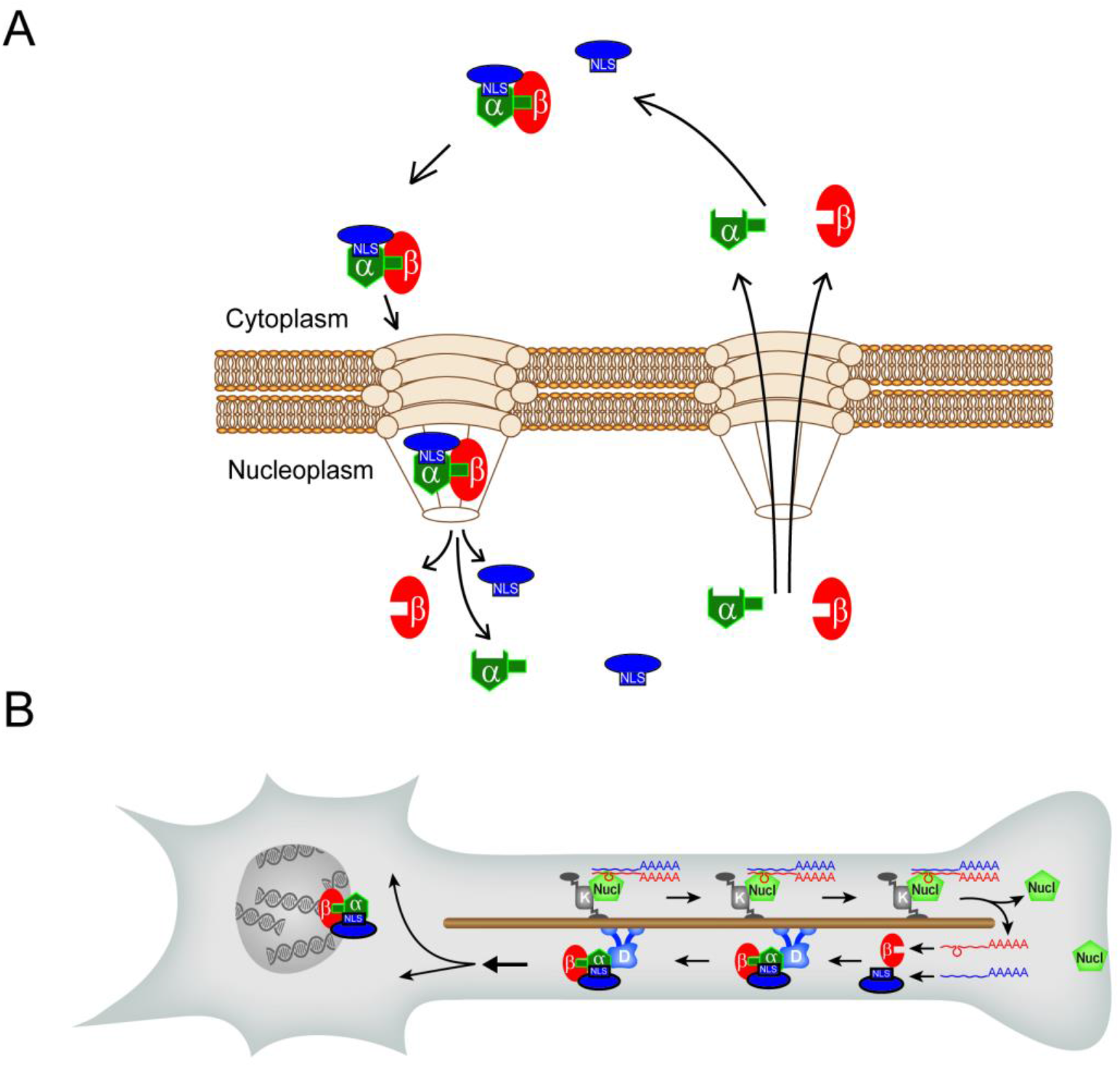
Importin β1 roles in intracellular transport. (A) Nucleocytoplasmic transport by importin β1 in complex with an importin α traffics NLS bearing cargos through the nuclear pore complex (NPC). (B) Importin β1 mediated transport in neuronal cytoplasm occurs upon local translation of anterogradely transported importin β1 mRNA, followed by formation of a retrogradely transported importins-cargo complex. K, kinesin; Nucl, nucleolin; D, dynein, β, importin β; α, importin α.

Subcellular localization is an important determinant of biological function, hence there is keen interest in determining importin interactomes and importin-dependent protein partitioning within cells (5). Comprehensive characterization of importin interactomes is challenging, due to the transient nature of importin-cargo interactions and the overlapping specificities of different importin complexes. Hence, most previous studies have focused on identifying importin carriers for specific cargo proteins, revealing a diverse range of both direct and indirect KPNB1 interactors (e.g. (6–10)). Other studies have attempted to identify importin cargos from transport perturbation studies (11, 12) or from transcriptome analyses after genetic perturbations (e.g. (13–16)). Unbiased proteome-wide studies have used proximity biotinylation to characterize an importin β1 interactome in HEK cells (17) and affinity proteomics to identify interactors in mitotic cells (18, 19). These diverse studies have unveiled an expanding list of KPNB1 interactors, but for the most part do not shed light on subcellular specializations in importin β1 interactions.

As noted above, importin β1 has specific cytoplasmic roles in cell size sensing and in neuronal injury signaling that are distinct from its nucleocytoplasmic transport activity (20–23). We therefore sought methods for comprehensive subcellular studies of its interactomes. A diverse range of proximity based biotinylation assays provide powerful tools to label and subsequently detect interacting proteins (24). Initial tests of BioID (25) suggested limited efficiency of the method for both N and C terminal fusions of KPNB1 with the modified biotin ligase BirA. We therefore generated a monoclonal antibody to enable biotinylation by antibody recognition (BAR) (26) for importin β1. The antibody targets a 20 amino acid epitope that is highly conserved in mammalian KPNB1 and discriminates between nuclear and cytoplasmic protein complexes, allowing a rigorous characterization of cytoplasmic importin β1 interactomes using the BAR method.

### Experimental Procedures

#### Animals

All animal procedures were performed in accordance with the guidelines of the Weizmann Institute of Science Institutional Animal Care and Use Committee (IACUC). Animals were purchased from Envigo Ltd (Israel) or originate from in-house breeding. Female BALB/c mice (6-8 weeks, Envigo) were used for immunization and subsequent isolation of spleen cells. Follow-up antibody characterization experiments were performed on male C57BL6 mice (8 - 12 weeks, Envigo) and on importin β1 3’UTR KO mice (8-12 weeks, in-house breeding (21)), as well as male Wistar rats (8-12 weeks, Envigo). All animals were maintained at the veterinary resources department of the Weizmann Institute with free access to food and water.

#### Sciatic nerve crush

The animals were anesthetized with ketamine/xylazine (10 mg / kg of body weight intraperitoneal). The sciatic nerve (SN) crush was performed at the mid-thigh level using fine jeweler forceps in two adjacent positions for 30 seconds each. Only one side was subjected to the crush injury, the contralateral side served as an uninjured control. Sciatic nerves were excised 6 and 24 hours after injury, fixed and processed for immunohistochemistry.

#### Reagents and antibodies

Commercially available primary antibodies (ABs) used in this study for immunoprecipitation (IP), immunofluorescence (IF), proximity ligation assay (PLA), Western blot (WB) and Simple Western capillary system (WES) were: Anti-GFP (abcam, ab6556, IF & WB 1:5000, PLA 1:2000), anti-NFH (abcam, ab72996, 1:2000), anti-Actin (MP, clone C4, 08691001, WB 1:5000), anti-GAPDH (Santa Cruz Biotechnology, sc-32233, WB 1:5000), anti-ERK1/2 (Sigma, M5670, WB 1:30000), anti-β-III tubulin TUJ1 (abcam, ab18207, IF 1:1000, WB: 1:6000), anti-Dynein heavy chain (DYNC1H1, Proteintech, #12345-1-AP, PLA 1:50), anti-importin beta 1 (MBS713065, rabbit, polyclonal, 1:1000).

#### Plasmids and constructs

Expression constructs for KPNB1 were based on pEGFP-N1 human full length KPNB1 from Addgene (#106941). The KPNB1 CDS was amplified from pEGFP-N1 human full length KPNB1 (Addgene #106941) and mVenus CDS was amplified from pTrix-mVenus-PA-Rac1, Addgene #22007). For seamless protein engineering of importin β1 BioID constructs, all cloning steps were carried out using restriction-free (RF) cloning following published protocols (27, 28). All primers were designed such that the protein domains (BirA*, ImpB1 and mVenus) are separated by a small flexible GS-dipeptide. Briefly, the MCS of pcDNA3.1 mycBioID (Addgene #35700) was removed to generate the Myc-BirA* control. Subsequently, the KPNB1 coding sequence was inserted either downstream or upstream of BirA* to generate N- or C-terminally BirA*-tagged importin β1 fusion constructs respectively. The opposite terminus of importin β1 was fused to mVenus (amplified from pTrix-mVenus-PA-Rac1, Addgene #22007) which has a similar size to BirA* to ensure similar molecular behavior of both fusion constructs. Detailed cloning procedure described in the supplementary data.

#### Generation of Monoclonal Antibodies

The antibody was raised against full-length recombinant mouse importin β1 protein (UniProt accession # P70168) with an N-terminal His-tag, produced in yeast (MyBioSource: cat # MBS955236). Importin β1 protein was dissolved in 1xPBS and mixed with Complete Freund’s Adjuvant (CFA) in a double-glass syringe at a volume ratio of 1:1 (23 µg protein in 50 µl 1xPBS +50 µl CFA / mouse) for immunization. Five BALB/c female mice (6-8 weeks old) were used for immunization by footpad injection into both hindlimbs (50 µl/food pad). Two weeks later, mice were boosted with the same dose (boost 1: 23 µg protein in 100 µl / mouse) intradermally in several spots in the belly area and limbs (close to lymph nodes). Retroorbital bleeding was performed 1 week later and serum was tested by ELISA (enzyme-linked immunosorbent assay) to determine the mouse with the highest immune response. Four weeks after boost 1, the mouse with the highest immune response received a prefusion boost (IP injection of 20 µg protein in 200µl 1xPBS, without CFA) and was sacrificed four days later to collect immune cells from the spleen. ∼100 million spleen cells were harvested and collected in DMEM for the production of hybridoma cell lines. The timeline of the immunization procedure is shown in Figure S2A.

Hybridomas were generated as previously described (29). BALB/c female mouse splenocytes were fused with NS0 non-secreting murine myeloma cells using polyethyleneglycol 41% in DMEM (PEG, Roth). Hybrid cells were selected in growth medium supplemented with HAT (hypoxanthine, aminopterin, and thymidine) (100x HAT media supplement, Sigma-Aldrich). Culture supernatants from wells with viable clones were screened by an indirect ELISA using recombinant importin β1 protein. Stable hybridoma clones secreting importin β1 specific antibodies were obtained after two cloning cycles by a limiting dilution assay and were weaned from HAT supplements. Hybridoma cells were grown in complete Dulbecco’s modified Eagle’s medium (DMEM, Biochrom) supplemented with 15% fetal bovine serum (FBS, Biochrom), 2mM L-glutamine, 10,000 u penicillin, 10,000 u streptomycin, 1mM Na pyruvate, with or without HAT supplements (1:100 from stock). The final positive hybridoma cells secreting antibodies against importin β1 were stored in liquid nitrogen. All procedures involving experimental mice were performed under controlled laboratory conditions in strict accordance with IACUC guidelines. A schematic of the selection process and hybridoma propagation is shown in Figure S2B. For more detailed protocols see supplementary material.

#### Enzyme-linked immunosorbent assay (ELISA)

Indirect ELISA was performed in 96-well plates (NUNC-442404), coated with full length importin β1 protein (1µg/ml in 1xPBS) overnight at 4°C and then blocked by incubation with 10 mg/ml BSA in 1xPBS for 1h at RT. Hybridoma supernatant or purified and diluted Abs were applied and incubated for 1h at RT, followed by HRP-conjugated anti-mouse Fab fragments (1h RT, 1:60000) as the secondary antibody (Sigma A9917-1ml). HRP substrate (TMB Sigma T0565-100ml) was added subsequently and the absorbance was recorded at 630 nm using a microplate reader (Thermo Fisher Scientific). In between steps, plates were washed in PBST (PBS containing 0.05% Tween-20) three times. For linear ELISA coating was done with a serial dilution of the full length importin β1 protein (0.001, 0.01, 0.1, 1 µg/ml), to explore the linear range of antibody-antigen recognition.

#### Antibody isotyping & purification

Immunoglobulin isotyping test was performed using a mouse monoclonal antibody isotyping kit, according to the manufacturer’s instructions (SBA Clonotyping System-HRP, SouthernBiotech, 5300-05). Monoclonal antibodies were purified by Protein A affinity purification, using an AKTA PURE system. The purified fraction was desalted in 1XPBS, using a HiPrep desalting column, and stored at 2-8°C.

#### DRG culture

Adult mouse dorsal root ganglia (DRG) neuron culture was performed as previously described (30). Briefly, the ganglia were dissociated with 100 U of papain (P4762, Sigma) followed by 1 mg/ml collagenase-II (11179179001, Roche) and 1.2 mg/ml dispase-II (04942078001, Roche). The DRGs were triturated in 1xHBSS, 10 mM glucose, and 5 mM HEPES (pH 7.35) using a fire-coated Pasteur pipette and then layered on 20 % percoll in L15 media and recovered by centrifugation at 1000 g for eight minutes. Cells were washed briefly in growth media (F12, 10 % Fetal Bovine Serum (FBS), Primocin (100µg/ml, InvivoGen #ant-pm-1)) and plated on poly-L-lysine (P4832, Sigma) and laminin (23017-015, Invitrogen) coated glass cover slips. Culture media and serum were purchased from Thermo Fisher Scientific. DRG neurons were grown for 24-48h in culture at 37°C and 5% CO_2_.

#### Cell line cultures

HeLa cells (human, female, RRID: CVCL-0030), HEK-293 (human, female, RRID: CVCL_0045), NIH/3T3 (mouse, male, RRID: CVCL_0594) and Neuro-2A (mouse, male, RRID: CVCL_0470) cells were purchased from ATCC (Cat# CCL-2,CRL-1573, CRL-1658, CCL-131, respectively). All cell lines were cultured in DMEM (Gibco), supplemented with 10% fetal bovine serum (Gibco), 100 U/mL *penicillin* and 100 μg/mL streptomycin. All cells were incubated at 37°C and 5% CO_2_.

#### Protein extraction for SDS Page, Western Blot, and WES

For total protein extraction, tissues were collected in RIPA buffer (150 mM NaCl, 1.0 % NP-40, 0.5 % sodium deoxycholate, 0.1 % SDS, 50 mM Tris, pH 8.0.), supplemented with protease/phosphatase/RNase inhibitors (complete protease inhibitor EDTA free (Roche 1187358000), phosphatase inhibitor cocktail 2 (1/1000, Sigma 5726), phosphatase inhibitor cocktail 3 (1/1000, Sigma P0044) and RNAse inhibitor (200U/ml, RNAseIn, Promega N251B)). Tissue samples were homogenized in Dounce Tissue Grinders (WHEATON 33, 1 ml #357538, 7 ml #357542) or, using plastic pestles, in an Eppendorf tube.

The axoplasm for biochemical analysis was extracted from mouse, rat, or bovine SN as previously described (31). To minimize glia contamination, transport buffer (TB) was used in this extraction protocol (20 mM Hepes, 110 mM KAc, 5 mM MgAc, pH 7.4 supplemented with protease/phosphatase/RNase inhibitors). All protein extracts were incubated on ice for 20 min, followed by a spin at 10,000 x g for 10 min at 4°C.

For Western blot, proteins were blotted on nitrocellulose membranes using a Trans-Blot Turbo™ Transfer System (Bio-Rad). Membranes were blocked with 5% nonfat dry milk in 1xTBST buffer for one hour. Primary antibodies were incubated for one hour at room temperature or overnight at 4°C with shaking. Secondary HRP-conjugated antibodies were diluted 1:10,000 in 1xTBST and incubated for 1 hour at room temperature. Blots were developed using Radiance ECL substrate (Azure biosystems) or SuperSignal™ West Femto (Thermo Fisher Scientific) substrates on an Amersham Imager 680. Automated capillary electrophoresis immuno-quantification runs were conducted on a WES instrument (ProteinSimple) as described (32). The indicated MW are based on an internal standard spiked into every capillary.

#### Proximity ligation assay (PLA)

PLA was performed on cultured DRG neurons using the Duolink system (Sigma) according to the manufacturer’s instructions with minor modifications. DRG neurons (isolated from C57BL6 mice) were fixed in 4% PFA after 48 hours. in culture. Blocking was performed in 5% donkey serum with 1% BSA in 1xPBS for 30 min, followed by primary antibody incubation for one hour at room temperature. Combinations: rabbit anti-DYNC1H1 (Proteintech, #12345-1-AP, 1:50) or rabbit anti-importin β1 (MBS713065, 1:1000), together with the antibody generated in the current study mouse anti-importin β1 (mAbKPNB1-301-320, 1 μg/ml). As controls, each antibody was used by itself and furthermore, we used mAbKPNB1-301-320 in combination with another rabbit antibody (GFP, abcam, ab6556, 1:2000). Interactions (PLA signals) were detected using Sigma PLA probe anti-mouse minus DUO92004, anti-rabbit plus DUO92002, and detection kits Red DUO92008 or FarRed DUO92013 according to the manufacturer’s instructions and incubation times. The PLA protocol was followed by IF staining for NFH (abcam, ab72996, 1:2000, 45 min at RT) with Alexa 488 donkey anti-chicken secondary antibody (45 min. at RT). The quantification of the PLA signal was performed on NFH positive axons using Fiji (33).

#### Electron Microscopy

DRG neurons were grown on sapphire disks and fixed 48 hrs. after plating using high-pressure freezing in a Bal-Tec HPM10, followed by freeze substitution, washing, embedding in HM20, and ultrathin sectioning (70–90 nm). Sections were collected on nickel grids. For immunostaining, grids were reacted with different concentrations of mAbKPNB1-301-320 (0.7mg/ml stock, dilutions: 1:2, 1:5, 1:10), followed by secondary anti-mouse immunoglobulin G with 10μm gold particles. The grids were then stained in uranyl acetate and lead citrate and analyzed under 120 kV on a Tecnai 12 (FEI) transmission electron microscope with an EAGLE (FEI) CCD camera using TIA software.

#### Pull-down from rat brain

Whole brain was isolated from adult Wistar rats (male, 8-12 weeks, Envigo) and homogenized in 7 ml Dounce Tissue Grinders (WHEATON 33, 7 ml #357542) in transport buffer, supplemented with NP40 (20 mM Hepes, 110 mM KAc, 5 mM MgAc, pH 7.4 supplemented with complete protease inhibitor, phosphatase inhibitor cocktail 2 & 3, as well as 0.5% NP40 for the initial lysis, then for the IP diluted to 0.035%). For each IP sample, 1 mg of total brain extract was used and adjusted with TB to a total volume of 300µl and a final concentration of 0.035% NP40. 10% (30µl) was taken as input. The remaining lysate for the IP was incubated for 3h at 4°C with overhead rotation with 10µg of mAbKPNB1-301-320 or control (pre-blocked mAbKPNB1-301-320: 10µg Ab with 50µg blocking peptide (KPNB1_301-320_) for 1h). Subsequently, 100 μl Protein G magnetic beads (Dynabeads, Thermo Fisher Scientific 10004D), pre-blocked with salmon sperm DNA (10μl DNA per 100μl beads for 1 hour 4°C), were added to the brain lysate - Ab mixture and incubated for additional two hours at 4°C. Beads were washed in several steps with different buffers as follows: TB-0.1% NP40 for three min., TB-0.5% NP40 three min., TB-1% NP40 one min., TB- no detergent three minutes. For simple western, elution from beads was conducted by denaturing the proteins from the beads with WES sample buffer supplemented with DTT to a final concentration of 40mM for five minutes at 95°C.

#### BioID in HeLa cells

HeLa cells were transfected using jetPEI® (Polyplus-transfection®SA) according to the manufacturer’s protocol and supplemented with 50 μM Biotin (Molecular Probes) for 24 hours. Cells were grown on 10 cm plates for mass spectrometry analyses. For streptavidin-IP from whole cell lysates, cells were lysed with 1.5 ml/plate Lysis buffer (50 mM Tris HCl, pH 7.5, 150 mM NaCl, 0.25% (w/v) NP-40, 1 mM MgCl_2_, 5 mM KCl, 2.5 mM EDTA, 1% (w/v) Triton X-100, 0.1% (w/v) SDS, 1 mM DTT, 1 x Complete protease inhibitor, EDTA-free (Roche)), sonicated (30% amplitude, 10 cycles of 5.5 sec pulse on, 9.9 sec pulse off) and lysates from six plates were incubated with 600 μl streptavidin beads for four hours. Beads (Dynabeads® MyOne^TM^ streptavidin C1, ThermoFisher Scientific) were then washed twice with Wash buffer I (2% (w/v) SDS,), once with Wash buffer II (0.1% (w/v) deoxycholic acid, 1% (w/v) Triton X-100, 1mM EDTA, 500 mM NaCl, 50 mM Tris HCl, pH 7.5), once with Wash buffer III (0.5% (w/v) deoxycholic acid, 0.5% (w/v) NP-40, 1mM EDTA, 250 mM LiCl, 10 mM Tris HCl, pH 7.5), twice with 50 mM Tris HCl, pH 7.5 and once with (20 mM Tris HCl pH 8.0, 2 mM CaCl_2_) (34) prior to processing for mass spectrometry.

#### BAR assays in HEK cells and DRG neurons

HEK 293T (human embryonic kidney) cells were grown in Dulbecco’s modified Eagle’s medium (DMEM) high glucose 5g/l (Sigma-Aldrich) supplemented with 10% heat inactivated fetal bovine serum (FBS). Cells were seeded at a density of 1.2e4 cells/cm^2^ and monitored over 24 hours until reaching 70% confluency. Cells were then fixed in 4% PFA and permeabilized with TritonX-100 for 10 minutes at RT, subsequently, incubated for three hours at 4°C with blocking solution. 2.7µg/ml mAbKPNB1-301-320 was added in blocking buffer and incubated over night at 4°C. The next day, cells were washed three times with 1xPBST and incubated with secondary HRP conjugated antibody for three hours at 4°C prior to TSA amplification according to the manufacturer’s protocol. Cells were harvested and boiled for one hour at 99°C for reverse crosslinking followed by streptavidin beads affinity purification and processing for proteolytic digest and mass spectrometry.

Adult DRG neuronal cultures were prepared as previously described (30). The culture was maintained for 72 hours with daily supplement of glia inhibitor AraC (10μM). The cells were then fixed in 4% PFA for 20 min at room temperature followed by three washes with 1xPBST, permeabilization with 0.1% TritonX-100 and incubation in blocking solution for three hours at 4°C. 2.7µg/ml mAbKPNB1-301-320 was added in blocking buffer and incubated over night at 4°C. The next day, cells were washed three times with 1xPBST and incubated with secondary HRP conjugated antibody for 3 hours at 4°C. The TSA amplification was performed according to the manufacturer’s protocol. The cells were then harvested and boiled for 1 hour at 99°C for reverse crosslinking followed by purification of the streptavidin beads affinity before processing for mass spectrometry.

#### Mass spectrometry-analysis for BioID assay samples

Sample-incubated streptavidin magnetic beads were resuspended in 20µl 5 mM DTT 100mM NH_4_HCO_3_ and incubated for 30 min at room temperature. Subsequently, iodoacetamide was added to a final concentration of 7.5 mM and the samples were incubated for 30 additional minutes. 0.5µg of sequencing grade trypsin (Promega) was added to each sample and incubated at 37C overnight. Supernatants of the beads were recovered, and beads digested again using 0.5ug trypsin in 100mM NH_4_HCO_3_ for two hours. Peptides from both consecutive digestions were combined and recovered by solid phase extraction using C18 ZipTips (Millipore), eluted in 15µl 50% acetonitrile 0.1% formic acid, and resuspended in 5µl 0.1% formic acid for analysis by LC-MS/MS.

Peptides resulting from trypsinization were analyzed on a QExactive Plus (Thermo Scientific), connected to a NanoAcquity™ Ultra Performance UPLC system (Waters). A 15-cm EasySpray C18 column (Thermo Scientific) was used to resolve peptides (90-min 2-30% gradient with 0.1% formic acid in water as mobile phase A and 0.1% formic acid in acetonitrile as mobile phase B. Mass spectrometer was operated in positive ion mode and in data-dependent mode to automatically switch between MS and MS/MS. MS spectra were acquired between 350 and 1500 m/z with a resolution of 70000. The top 10 precursor ions with a charge state of 2+ or higher over the selected threshold (1.7E4) were fragmented by HCD using a normalized collision energy of 25 with an isolation window of 4 m/z. MSMS spectra were acquired in centroid mode with resolution 17500 from m/z=100. A dynamic exclusion window was applied which prevented the same m/z from being selected for 10s after its acquisition.

For peptide and protein identification, peak lists were generated using PAVA in-house software (35). All generated peak lists were searched against the human subset of the SwissProt.2019.07.31 database (containing 20432 entries) using Protein Prospector (36) with the following parameters: Enzyme specificity was set as Trypsin, and up to two missed cleavages per peptide were allowed. Carbamidomethylation of cysteine residues was allowed as fixed modification. N-acetylation of the N-terminus of the protein, loss of protein N-terminal methionine, pyroglutamate formation from of peptide N-terminal glutamines, and oxidation of methionine were allowed as variable modifications. Mass tolerance was 10 ppm in MS and 30 ppm in MS/MS. The false positive rate was estimated by searching the data using a concatenated database which contains the original SwissProt database, as well as a version of each original entry where the sequence has been randomized. A 1% FDR was permitted at the protein and peptide level. All spectra identified as matches to peptides of a given protein (Peptide Spectral Matches, PSMs) were reported, and this number used for label free quantitation of relative protein abundances in the samples as ratios of each protein PSMs to the average PSMs for that protein on the appropriate control samples (cells transfected with Myc-BirA). Absent values were given a PSM value of 0.5. Statistical analysis was performed on three independent biological repeats using GraphPad Prism software applying multiple unpaired t test with a desired FDR of 10%.

#### Mass spectrometry-analysis of BAR assay samples

On bead digestion was performed as indicated above. Peptides resulting from trypsinization were analyzed on an Orbitrap Lumos Fusion (Thermo Scientific), connected to a NanoAcquity™ Ultra Performance UPLC system (Waters). A 15-cm EasySpray C18 column (Thermo Scientific) was used to separate peptides in a 60-min 2-30% gradient with 0.1% formic acid in water as mobile phase A and 0.1% formic acid in acetonitrile as mobile phase B. MS was operated in positive ion mode and in data-dependent mode to automatically switch between MS and MS/MS. MS spectra were acquired between 375 and 1500 m/z with a resolution of 120000. For each MS spectrum, multiply charged ions over the selected threshold (2e4) were selected for MSMS in cycles of 3 seconds, with an isolation window of 1.6 Th. Precursor ions were fragmented by HCD using a normalized collision energy of 30. MSMS spectra were acquired in centroid mode with resolution 30000 from m/z=110. A dynamic exclusion window was applied which prevented the same m/z from being selected for 30s after its acquisition MS data analysis was performed as indicated for BioID samples above, but in the case of data form DRG mouse neurons the mouse subset (17009 entries) of the SwissProt.2019.07.31 database was used to search the data. Statistical analysis was performed on three independent biological repeats for DRG samples and two biological repeats for HEK samples using GraphPad Prism software applying multiple unpaired t test with a desired FDR of 10%. The mass spectrometry proteomics data have been deposited to the ProteomeXchange Consortium via the PRIDE (37) partner repository with the dataset identifier PXD032728.

#### Immunofluorescence and Immunohistochemistry on Cryosections

Sensory neurons were grown on glass coverslips coated with Poly-L-Lysin and Laminin for two days and then fixed using 4% PFA, permeabilized with 0.2% Triton X for 10 minuts at 25°C and blocked for one hour at room temperature with 5% BSA in 1XPBST (1xPBS plus 0.1% Tween-20). Coverslips with neurons were incubated in mAbKPNB1-301-320 with or without blocking peptide KPNB1_301-320_, with a ratio of 4.6nM mAbKPNB1-301-320 to 9.5nM peptide, overnight at 4°C. After three five-minute washes in 1xPBS, cells were incubated in donkey anti-mouse FITC secondary antibodies (1:200 each; Jackson Immuno-Res) for one hour at room temperature. After three 5 minute washes in 1xPBS, coverslips were rinsed in ddH_2_O then mounted with Prolong Gold Antifade with DAPI.

Sciatic nerves of adult WT and 3’UTR importin beta 1 KO mice were dissected at the designated times after crush injury and non-injured controls, fixed in 4% PFA for 12 hours at 4°C. Then nerves were washed three times in 1XPBST (10 min each), incubated overnight in 30% sucrose and subsequently embedded in OCT freezing medium. The 12μm thick sections were cut with a cryostat and mounted on glass slides. Freshly cut sections were incubated in blocking solution (5%BSA in 1XPBS) for two hours at 4°C and incubated overnight with primary antibody (in 1XPBST with 2%BSA). Primary antibody solution was carefully removed and slides were washed three times (five minutes each) at room temperature with 1XPBST. In the next step the slides were incubated in donkey anti-mouse FITC secondary antibodies (1:1000 each; Jackson Immuno-Res) for two hours at room temperature. After three five-minute washes in 1XPBST, slides were mounted with Prolong Gold Antifade.

#### siRNA knockdown

N2A cells were seeded in six-well plates with 120,000 cells per well, one day before siRNA transfection. The cells were transfected with custom siRNA against the 3’UTR of mouse importin β1 (siKPNB1) or siControl #5, using DharmaFECT 4 transfection reagent (Dharmacon) according to the manufacturer’s protocol. 48 hours after transfection the cell growth medium was replaced by serum free medium to slow down proliferation and to activate the differentiation process. 72 hours after transfection the cells were harvested and total protein was extracted in RIPA buffer containing 0.1% SDS and protease inhibitors. 20 μg of total protein per each sample were run on gel for WB analysis.

#### Statistical methods

Data represent mean ± standard error of the mean (SEM), unless otherwise noted. Groupwise analyses were conducted by one- or two-way ANOVA with post hoc tests as described in the figure legends. Pairwise analyses were conducted by two-tailed Student’s t tests (unpaired, unless otherwise noted; see figure legends). Statistical analyses were conducted using GraphPad Prism, Synergy Kaleidagraph, or Microsoft Excel. Statistically significant p values are shown as * p < 0.05, ** p < 0.01, *** p < 0.001 and **** p < 0.0001. Physical subnetwork analysis was performed using String database version 11.5 (38). An interaction score of 0.4 was selected and the confidence based on all available active interaction sources.

Gene ontology analysis was done using metascape (39) to identify all statistically enriched terms, with a P value cutoff of 0.01 and min enrichment of 1.5. The significant terms were hierarchically clustered into a tree diagram, based on Kappa-statistical similarities among their gene memberships. A 0.3 kappa score was applied for term clustering. A selected a subset of representative terms from this cluster was converted into a network layout, in which each term is represented by a circle node, and its color represent its cluster identity. Terms with a similarity score > 0.3 are linked by an edge (the thickness of the edge represents the similarity score). The network was visualized with Cytoscape (v3.1.2) with a ‘force-directed’ layout and with edge bundled for clarity.

## Results

### BioID proximity labeling of importin β1 interactors in HeLa cells

We first tested BioID for proximity labeling of importin β1 interactomes. The effective labeling range of this method has been estimated as ∼10 nm (40), hence fusions of BirA* at the N-terminus of importin β1 might not label NLS containing cargos that bind the complex via importin α (Figure 2A, B). We therefore generated constructs with BirA* fused to either N- or C-terminus of the human importin β1 open reading frame, with a Myc epitope tag and Venus fluorescent protein at the opposite termini (Figure 2B). These and a control construct comprising only Myc-tagged BirA* were expressed in HeLa cells and expression was validated with either Myc or anti-importin β1 antibodies. Cells were pulsed with biotin for 24 hrs., and cell lysates were subjected to streptavidin-pull downs for mass-spectrometric analyses. A total of 277 specific hits were identified (Table S1), of which 54 hits, primarily nuclear pore complex proteins, were shared by both constructs (Figures 2C-E and S1). Additional candidates include proteins involved in cell division/spindle assembly, RNA splicing, mRNA metabolism and export from the nucleus, and proteins that participate in nuclear import (Figure S1). However, importin α’s were identified only with the C-terminal BirA* construct, and the dataset lacked many classical NLS-dependent importin cargos that are known to be expressed in HeLa cells. Since this might be due to the limited biotinylation range of BirA* (40), we generated a series of constructs incorporating linkers of different lengths between BirA* and importin β1. Despite extensive testing, none of these constructs revealed efficient tagging of co-transfected canonical importin α cargos, suggesting that steric hindrance and limited range may restrict the efficiency of importin β1 targeted BioID for tagging of endogenous importin cargos. Tagging an Importin α for BioID instead of importin β1 is not straightforward, since both the N- and C-termini of Importin α are essential for its functions (41–43). Biotinylation by Antibody Recognition (BAR) enables proximity labeling of interactors of a protein of interest upon antibody binding (26), and moreover has an effective radius of up to ∼0.5–1 µm (44, 45), which might be advantageous for importin interactome characterization. However, the approach is critically dependent on antibody specificity and sensitivity (26). We therefore set out to generate a high-quality monoclonal antibody for importin β1.

**Figure 2:**
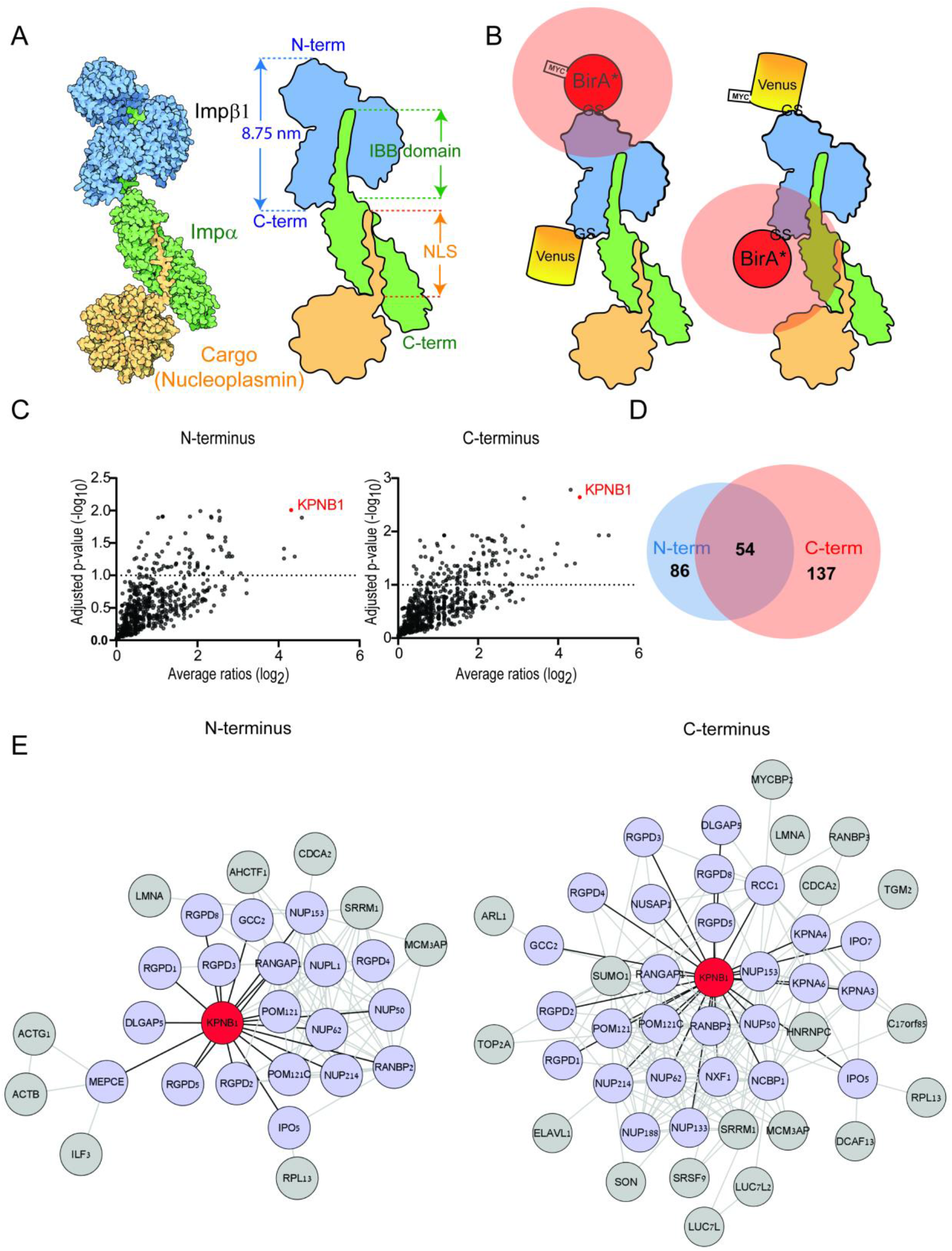
Importin β1 BioID. (A) Structural model of importins complex with an NLS-bearing cargo protein, nucleoplasmin, from https://pdb101.rcsb.org/motm/85. IBB, importin β1 Binding domain of importin α. (B) Schematic showing a set of KPNB1 BioID constructs superimposed on the structural model. Constructs have N- or C-terminal BirA* fusions, and Venus reporter with Myc tag on the opposing termini. GS, Glycine-Serine linker. Pink shading around BirA* indicates 10nm range of biotinylation. (C) Volcano plots of label free LC-MS/MS analysis from HeLa cells overexpressing N- or C-terminal BirA* KPNB1 constructs. (D) Venn diagram: numbers of significant hits obtained by each construct. (E) Physical network representation of N- and C-terminal KPNB1-BioID interactomes, based on STRING with a cut-off interaction score of 0.4.

#### Antibody Generation and Epitope Mapping

Full-length recombinant mouse importin β1 protein was used to immunize mice for monoclonal antibody (mAb) production. A conventional immunization protocol was conducted on five animals, and the highest responding mouse was selected after verification of serum titer and specificity (Figures S2 and S3). The selected mouse underwent a pre-fusion boost four days prior to spleen cells collection and hybridoma fusion using standard procedures (29). Cell culture supernatants from ∼ 960 hybridoma clones were screened for secretion of anti importin β1 antibodies, yielding 20 positive clones. Further screening of these 20 clones included WB for endogenous KPNB1 in bovine axoplasm (Figure S3C), immunofluorescence in different cell types (HeLa, 3T3, mouse DRG, Figure S3D) and ELISA, using linear dilution series (Figure S3E). Clones were scored on a scale from 0 (no signal) to 3 (highest signal) in all applications and the top eight clones underwent subcloning, antibody purification and isotyping (Figure S3F). Purified antibody from these eight clones were validated again by ELISA (Figure S3G). All these eight antibodies were able to detect importin β1, with some mainly showing nuclear signal, while others revealed a clear preference for cytoplasmic importin β1. The antibody with the highest overall score from those with preferential immunoreactivity for cytoplasmic importin β1 was chosen for further characterization.

We used a custom peptide library screen to identify the epitope recognized by the selected mAb. 58 overlapping peptides with a length of 20 amino acids (aa) each were synthesized to tile the entire amino acid (aa) sequence of murine importin β1, conjugated to N-terminal biotin, and screened in ELISA format on streptavidin-coated 96-well plates (Figure S4A). This procedure led to the identification of a 20 aa epitope sequence corresponding to residues 301-320 of human KPNB1, within HEAT repeats 7 and 8. The alignment of the mouse, rat, human and bovine importin β1 protein sequence shows complete homology in the epitope region (Figure 3A). We used PyMOL to visualize the identified epitope in the importin β1 structure (46, 47), as shown in Figure 3B. To validate the affinity of the mAb for the identified sequence, we conducted reciprocal linear ELISAs, using either varying peptide dilutions (Figure 3C) or varying mAb dilutions (Figure 3D) and observed a linear signal increase until saturation. The mAb effectively recognizes Importin β1 from different species in a number of applications, including ELISA, WB, IP (Figure S4), and Immunofluorescence (IF) (Figure S5). A direct comparison of the mAb performance in IF staining on several cell lines derived from different species is shown in Figure S5. We also tested whether the identified epitope peptide sequence can serve as a blocking agent in IP assays. Pre-incubation of the mAb with the peptide prior to IP from rat brain lysates, showed that KPNB1_301-320_ competes with binding of endogenous full-length importin β1, as demonstrated by reduced signal in the IP and no evident depletion in the supernatants (Figure S4). Based on the validated epitope identity, the antibody is henceforth termed mAbKPNB1-301-320.

**Figure 3:**
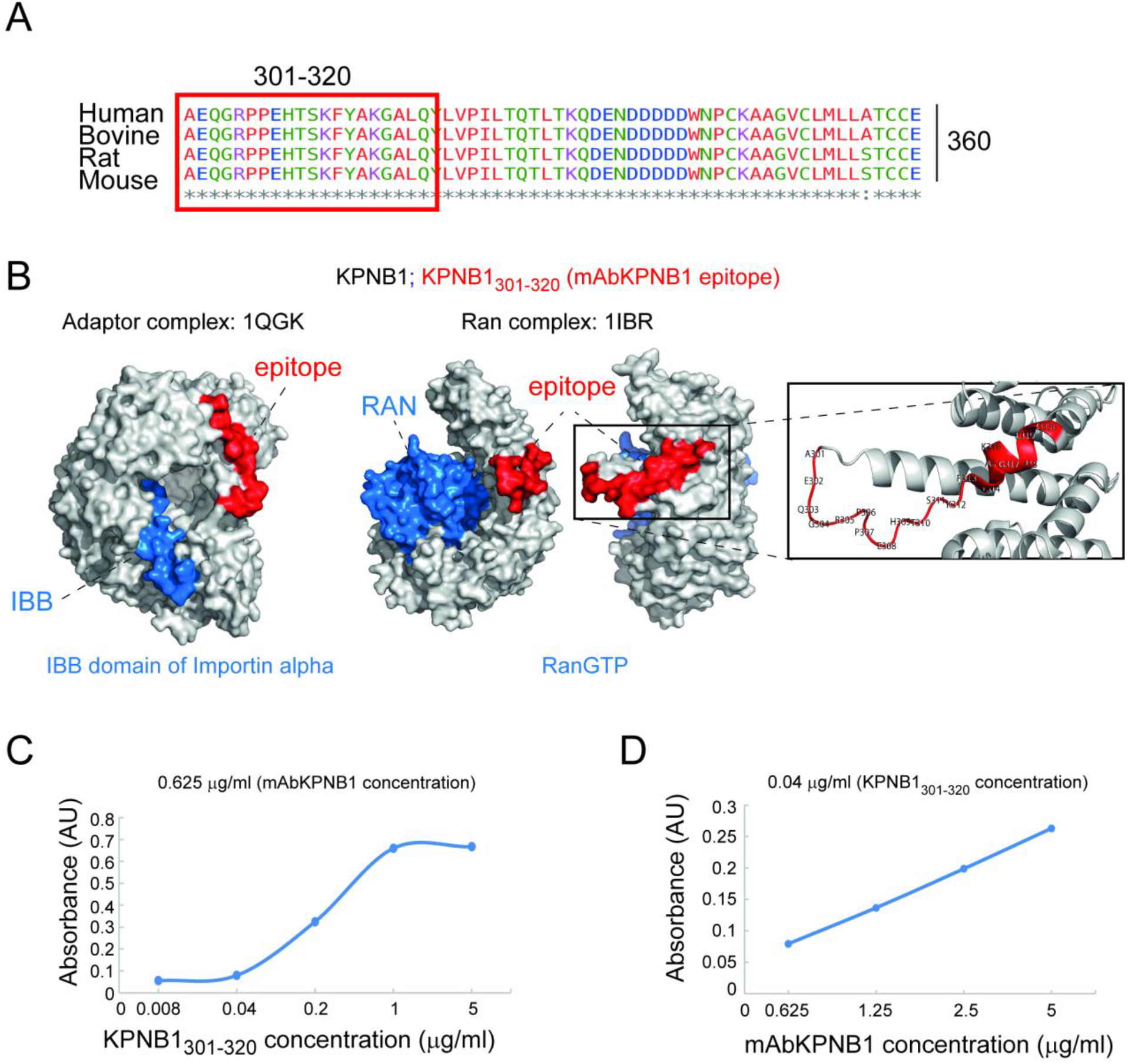
Epitope characterization of the mAb. (A) Multiple sequence alignment show homology of the identified epitope, comprising the residues 301-320 of importin β1, with color coding according to the physicochemical property of the amino acid by Clustal Omega. (B) PyMOL generated representations of importin β1 (grey) in complex with the N-terminal IBB domain of importin α (1QGK) or Ran (1IBR) (both in blue), with the identified epitope shown in red. Ran complex shown from two angles. (C-D) ELISA data with serial dilutions of either peptide (C) or mAb (D) confirms dose-dependent recognition.

#### mAbKPNB1-301-320 efficiently recognizes importin β1 in sensory neurons

Locally translated importin β1 is retrogradely transported from axon to soma in injured sensory neurons and is also associated with dynein in growing axons (20–22). We set out to test mAbKPNB1-301-320 in recognition of importin β1 in sensory axons. Indeed, immunostaining of cultured neurons revealed strong cytoplasmic staining, which was absent in cells incubated in parallel with a blocking peptide comprising the epitope sequence (Figure 4A, B). Recognition of axonal importin-β1 by mAbKPNB1-301-320 was also confirmed by immunogold labeling in EM sections of cultured sensory neurons (Figure 4C) and by in situ proximity ligation assay (PLA, Figure S6). The latter method allows the detection of two epitopes, either on the same protein or on two interacting proteins, within a radius of 0-40 nm (48). Here, we used mAbKPNB1-301-320 together with a commercially available polyclonal anti-KPNB1 to detect importin-β1 in axons of cultured DRG neurons. Furthermore, we used PLA to validate the ability of mAbKPNB1-301-320 to recognize importin β1 in association with dynein, robustly detecting colocalization of importin β1 and dynein heavy chain (Dync1h1) in axons of growing DRG neurons (Figure 4 D, E).

**Figure 4:**
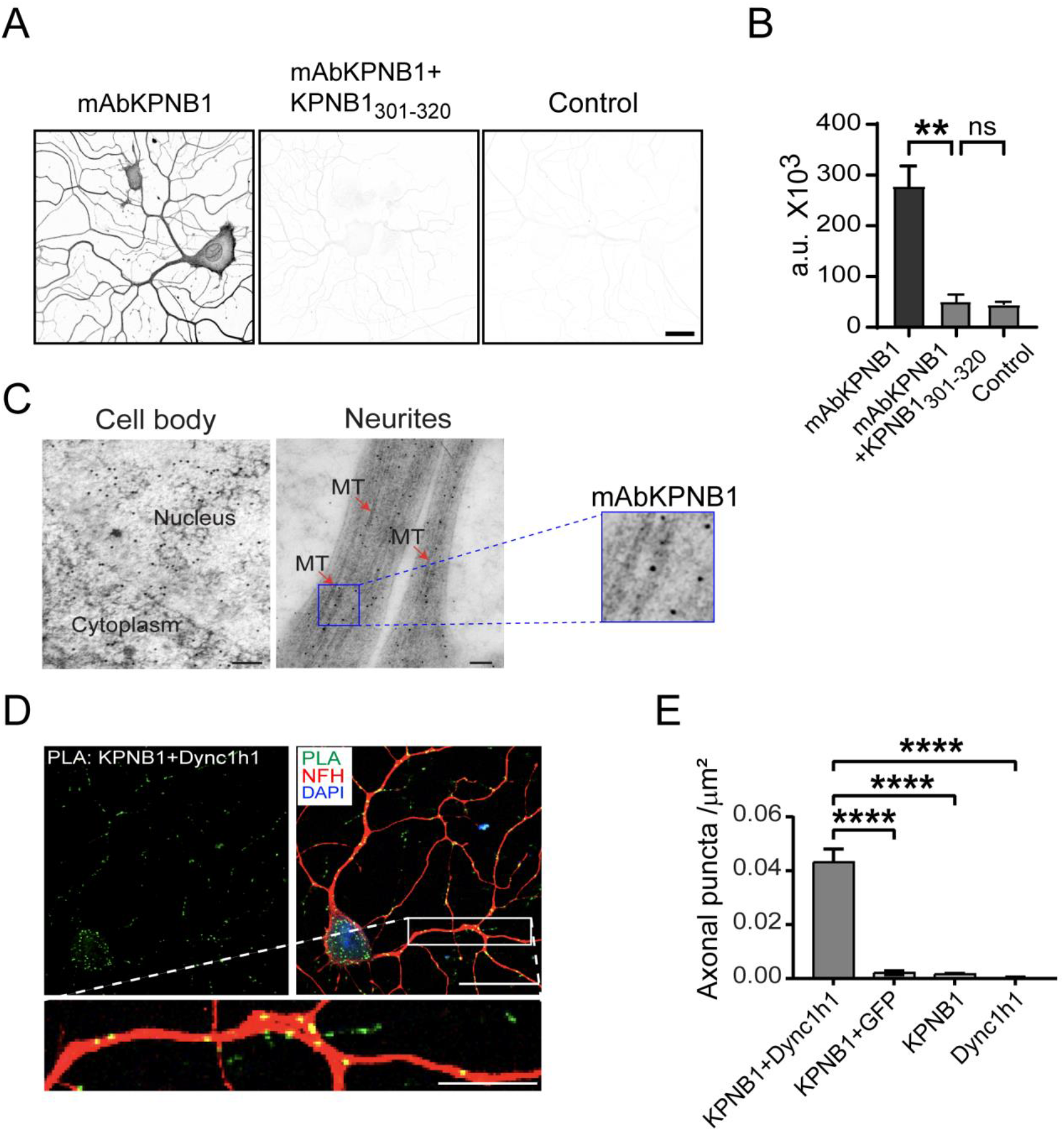
mAbKPNB1-301-320 recognizes importin β1 in adult DRG neurons. (A) Confocal images of adult DRG neurons in culture, stained for importin β1 using mAbKPNB1-301-320 (mAbKPNB1, 4.6 nM). Middle panel after pre-incubation with a peptide comprising the epitope sequence (9.5 nM KPNB1_301-320_), right panel is omitted primary antibody control. Scale bar 20 *¼*m. (B) Quantification of mAbKPNB1-301-320 average fluorescence intensity using ImageJ, Mean ± SEM, n≥5, one-way ANOVA, ** indicates *p* < 0.01. (C) Electron micrographs of ultrathin monolayer sections showing immuno-gold labeling of importin β1 in cultured mouse DRG neurons using mAbKPNB1-301-320 (mAbKPNB1, 0.23 μg/μl). Scale bars: 100 nm; gold particle diameter: 10 nm, MT: microtubules. (D) Proximity ligation assay (PLA, green) showing complexes of importin β1 (KPNB1) with dynein (Dync1h1) in axons of cultured DRG neurons. Neurofilament heavy chain (NFH, red) and DAPI (blue). Scale bars: full image 50 µm, insert 10 µm. (E) Quantification of PLA signal in axons, one-way ANOVA with Dunnett’s multiple comparisons test, **** indicates *p* <0.0001, n > 13.

In order to test the ability of mAbKPNB1-301-320 to detect axonal importin β1 upregulation *in vivo* we used the mouse model of sciatic nerve crush injury. Nerves were isolated six hours after crush, fixed and processed for immunohistochemistry (IHC) for frozen sections. mAbKPNB1-301-320 detected significant upregulation of importin β1 protein in injured sciatic nerve axons (Figure 5A, B). Local upregulation of importin β1 in axons after SN crush was also detected when using mAbKPNB1-301-320 for IHC on paraffin sections (Figure S7A, B). Furthermore, we used PLA to validate the ability of mAbKPNB1-301-320 to detect importin β1 in association with dynein in the sciatic nerve. PLA signal significantly increased after injury, confirming local translation of importin β1 protein upon nerve injury and it’s binding to dynein for retrograde transport (Figure 5C-E). Little or no background was detected in both injured and non-injured PLA probe and antibody controls (Figure S8).

**Figure 5:**
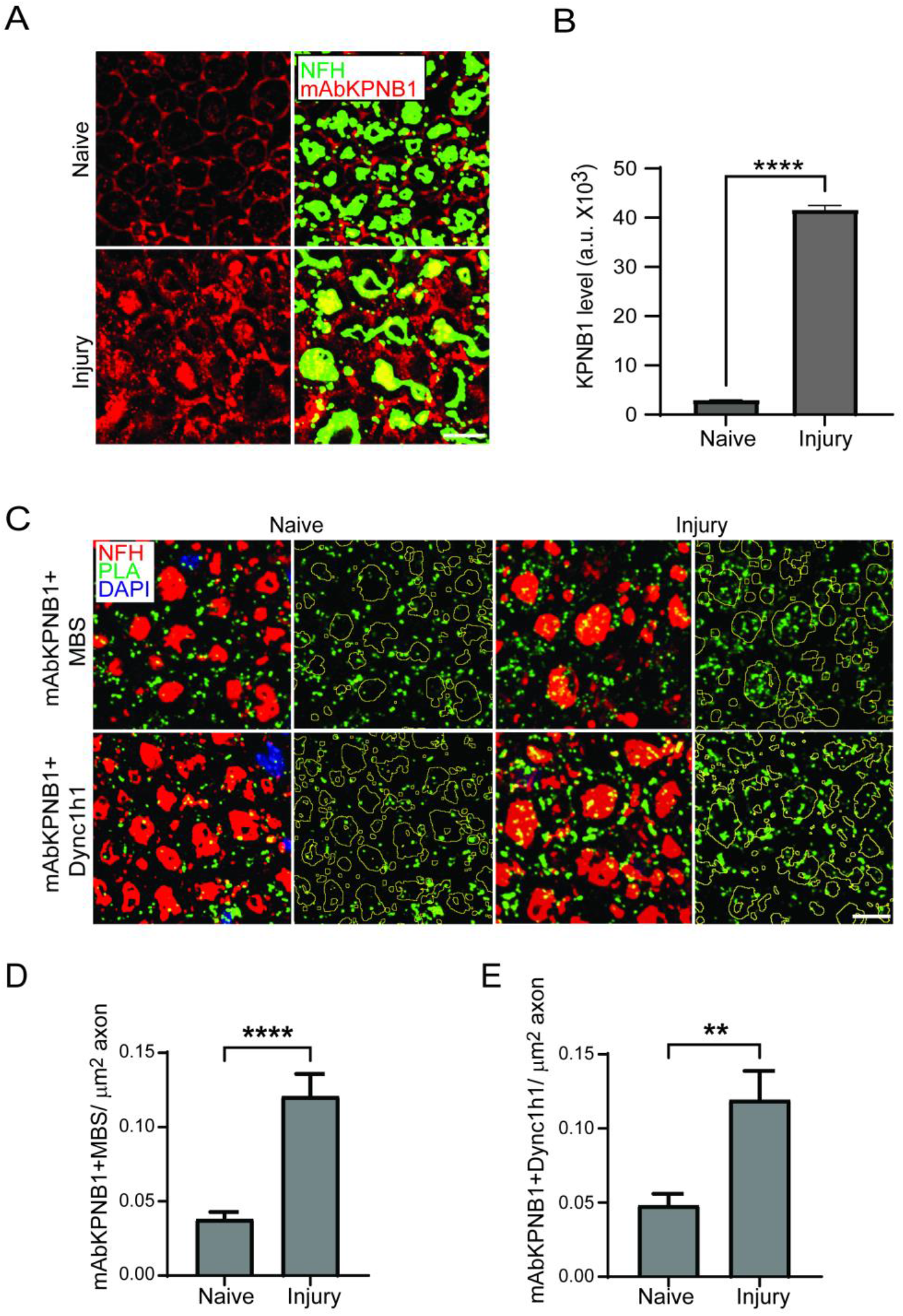
mAbKPNB1-301-320 detects importin β1 upregulation in injured nerve. (A) Immunostaining of cryosections with mAbKPNB1-301-320 (mAbKPNB1, 2.7ng/μl) before and 24 hrs. after sciatic nerve crush. Scale bar 10 μm. (B) Quantification of analysis shown in (A) in ImageJ, using NFH staining as a mask for axons. Mean ± SEM, n ≥ 490 axons, Unpaired two-tailed t test, **** indicates *p* < 0.0001. (C) Confocal imaging of PLA. Upper: PLA for importin β1 with mAbKPNB1 and a commercial anti-KPNB1 antibody (MBS); Lower: PLA for dynein and importin β1 with anti-Dync1h1 and mAbKPNB1; both on cross sections of the sciatic nerve, Scale bar 5 μm. (D, E) Quantification of the analyses shown in C, for importin β1 specific detection (D) or importin β1 in complex with dynein (E). Mean ± SEM, n > 2000 axons, Unpaired two-tailed t test, ** indicates *p* < 0.01, **** *p* < 0.0001.

#### mAbKPNB1-301-320 for BAR analyses of importin β1 cytoplasmic interactomes

After thorough validation of mAbKPNB1-301-320, we tested its utility in capturing importin β1 interactors using the BAR method on HEK cells (Figure S9). A total of 276 importin β1 proximal proteins were identified in this analysis (Figure S9). Candidate proteins include nucleocytoplasmic transport proteins and strictly cytoplasmic interactors, such as the translation initiation factor EIF4B and the stress granule protein G3BP1 (Figure S9C, D, Table S2). Comparison of this dataset to a previously published BioID dataset for KPNB1 in HEK cells (17) revealed approximately 80% novel candidates, with a bias for cytoplasmic components, as compared to more nucleoplasmic representation in the BioID dataset (Figure S9B). We then proceeded to use mAbKPNB1-301-320 for BAR analyses on DRG neurons grown for three days in culture. Streptavidin staining confirmed that biotinylation occurred in cytoplasm with negligible background signal from the nucleus (Figure 6A). A total of 432 specific candidates were identified in this experiment (Figure 6B, Table S3). String database analysis revealed that ∼28% of these proteins are annotated as of neuron projection origin and ∼18% as axonal proteins. Overall, 93% of the candidates were annotated as cytoplasmic proteins, some of these in overlap with hits annotated as occurring also in the nucleus. This overlap is not surprising for a nucleocytoplasmic transport protein. A number of functionally distinct complexes were identified, including components of the axonal growth cone, the dynein retrograde motor complex, axonal-dendritic transport proteins, Rho GTPase signaling systems, and markers for the cellular leading edge (Figures 6C, D, and S10). We also identified known importin β1 cargos such as ribosomal proteins, Stat3 and vimentin, Table S3 (6, 49, 50). These results confirm mAbKPNB1-301-320 as a high-quality probe for importin β1 interactomes.

**Figure 6:**
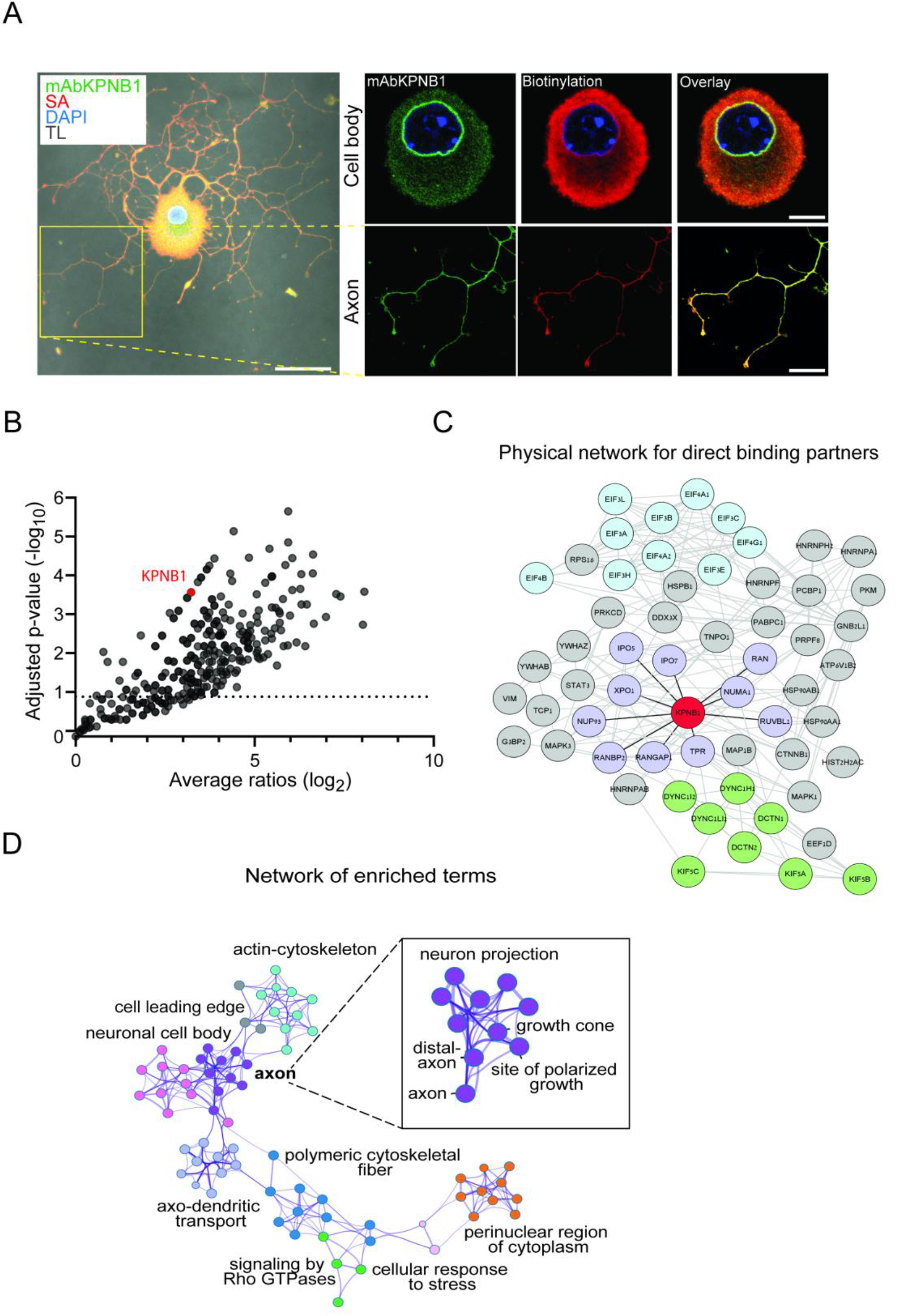
BAR characterization of the cytoplasmic importin β1 interactome in adult DRG neurons. (A) Visualization of mAbKPNB1-301-320 directed BAR reaction in a cultured DRG neuron, showing endogenous cytoplasmic importin β1 (green) and biotinylation (red). Scale bars: full image 40 μm, cell body 10 μm, axon 20 μm. Note the predominantly cytoplasmic and axonal biotinylation. (B) Volcano plot of proteins identified by mass spectrometry with BAR, multiple t-test with a desired false discovery rate of 10%, n=3. (C) Network analysis of hits from (B) showing subnetwork of direct binding partners based on STRING database. KPNB1 in red, retrograde motor interactors in green, translation machinery in light blue, nucleocytoplasmic transport in light purple. (D) Partial network of selected enriched terms with focus on the axonal compartment: colored by cluster ID, nodes that share the same cluster ID are closer to each other, generated by Metascape.

## Discussion

In this study, we generated and validated a new monoclonal antibody for importin β1 and demonstrated its utility in proximity biotinylation analyses of importin β1 interactomes using the BAR method. We turned to the BAR approach after initial attempts to use importin β1/BirA* fusions for BioID, which identified various nuclear and nucleocytoplasmic transport interactors but revealed poor efficiency for identifying cytoplasmic interactors of the importins complex. The chemistry on which the BAR method is based has a significantly larger labeling range than BioID (26, 45), and the method does not require the introduction of exogenous tags to the protein of interest. These characteristics are advantageous for comprehensive coverage of interactomes of interest, but conversely, they may decrease the specificity of the analysis. Hence, the effective use of BAR is critically dependent on the antibody quality. mAbKPNB1-301-320 was shown to be a highly specific and sensitive importin β1 probe in a broad range of applications, cell types and species. Mapping localized the epitope to amino acids 301-320 in importin β1, and importantly the linear 20-mer epitope peptide is effective as a blocking reagent in different assays. Visualization of the epitope in the crystal structure reveals a region in a concave part of importin β1 within the stalk motif (47). The epitope protrudes by ∼20 Å from the surface of the protein and is clearly accessible to the antibody in cytoplasmic and axonal compartments, but less available within nuclei. The significance of this subcellular specificity for importin β1 functions will be an intriguing topic for future exploration, and the antibody provides an excellent tool for this purpose.

Previous characterizations of importin β1 interactomes used a diversity of approaches, including BioID (17), SILAC-Tp (12, 51, 52), and co-immunoprecipitation methods (18, 19). Different cell types and biological paradigms were used in these studies, complicating direct comparison of those datasets with ours. However, gene ontology analyses allow more comprehensive comparisons, and these are shown for subcellular localization and canonical pathway analyses in Figure 7. Unsurprisingly, both BAR datasets from the current study showed a higher representation of cytoplasmic proteins than the previous datasets, reflecting the subcellular preference of the antibody. This subcellular specificity is also reflected in the signaling pathways identified by BAR versus previous BioID and pull-down approaches (Fig. 7A), highlighting the need for diverse approaches in multiple cell types for truly comprehensive profiling of such interactomes. It should however be noted that the BioID studies conducted with KPNB1 to date have all used first or second generation BirA* enzymes, and it is possible that enhanced activity or faster acting variants developed recently (24, 53) would be more efficient.

**Figure 7:**
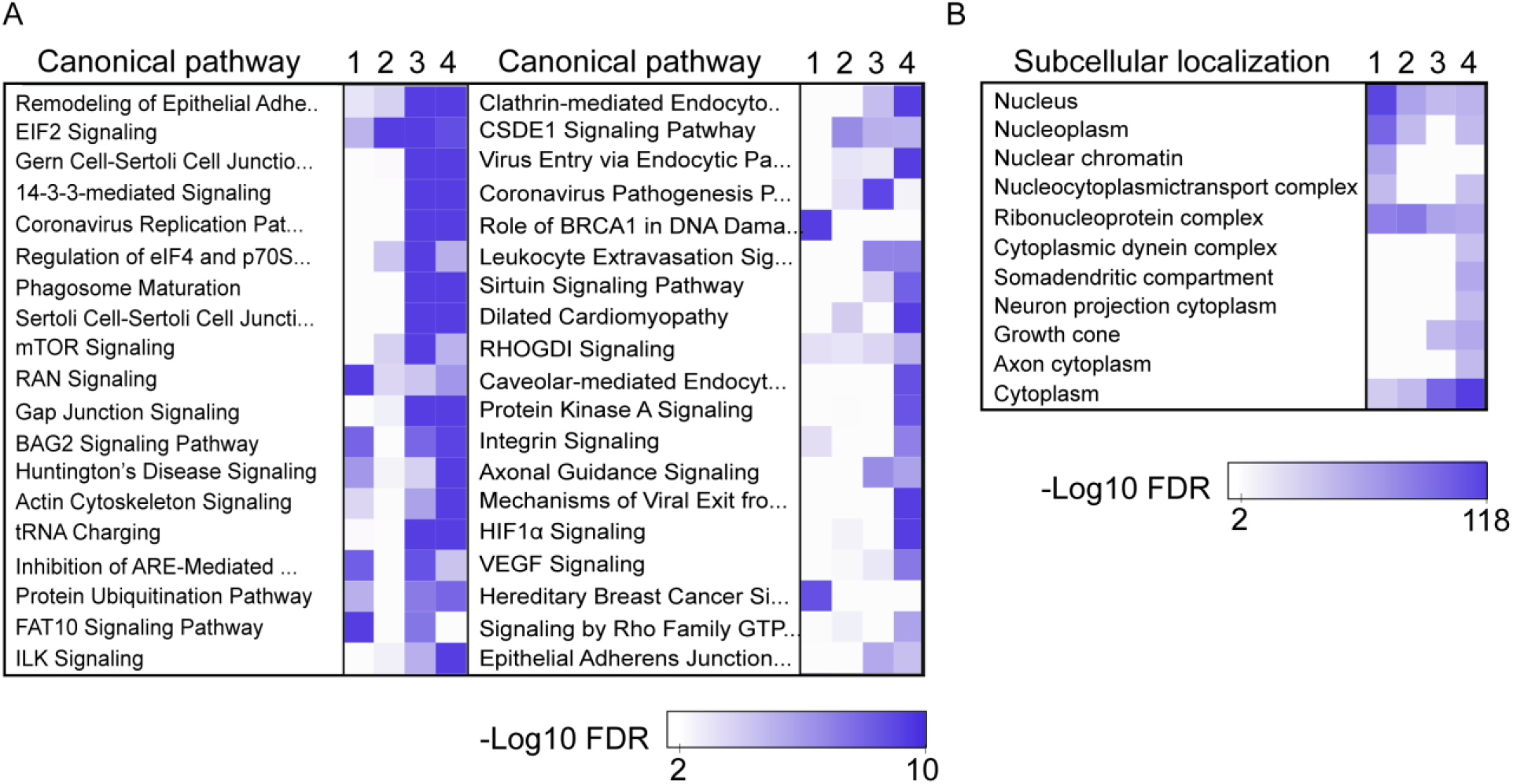
Gene ontology comparisons of different importin β1 interactomes. Gene ontology analyses from datasets of (1) Reference #17 (HEK293T), (2) References #18,#19 (HeLa) (3) HEK 293T BAR (4) DRG neuron BAR. (A) Canonical pathway comparison using Ingenuity pathway analysis sorted by score value and a 1% p value cut-off. (B) GO analysis using STRING categories for subcellular localization, p value cut-off: 1 %.

In summary, we have generated an antibody that discriminates between subcellular conformations of importin β1, and have shown that it provides a highly specific and sensitive probe for cytoplasmic importin complexes. BAR analyses utilizing the new antibody as a targeting agent reveal significant new categories of the importin β1 interactome, with focus on cytoplasmic and signaling proteins that highlight new functional roles of the importins complex that go beyond nucleocytoplasmic transport.

## Supporting information

Table S1 bioID HeLa list of hits

Table S2 BAR HEK list of hits

Table S3 BAR DRG list of hits

## Acknowledgements

We sincerely thank Prof. Zelig Eshhar and Tova Waks for their advice on monoclonal antibody generation; Vladimir Kiss, and Dr. Reinat Nevo for assistance with microscopy; Dr. Vera Shinder for assistance with electron microscopy; and Dr. Dalia Gordon for helpful discussions. We gratefully acknowledge funding from the Israel Science Foundation (ISF 1337/18 to MF & IR), and the Dr. Miriam and Sheldon G. Adelson Medical Research Foundation (to ALB & MF) and the Human Frontier Science Program Organization (to CA). MF is the incumbent of the Chaya Professorial Chair in Molecular Neuroscience at the Weizmann Institute of Science.

## Author contributions

D-AS, ED-M, SA, CA, MF, and IR designed the study; D-AS, SA, CA, ED-M, IR, VS, OL, and HH planned and performed experiments and data analyses; JAO-P carried out mass spectrometry analyses; MF, ALB, and IR supervised research; D-AS, SA, ED-M, MF, and IR wrote the manuscript draft. All authors revised the manuscript and approved the final version.

## Abbreviations

BAR: biotinylation by antibody recognition
BioID: proximity-dependent biotin identification
CDS: coding DNA sequence
DRG: dorsal root ganglia
ELISA: enzyme-linked immunosorbent assay
EM: electron microscopy
IF: immunofluorescence
IHC: immunohistochemistry
IP: immunoprecipitation;
KPNB1: karyopherin β1/importin β1
mAb: monoclonal antibody
NFH: neurofilament heavy chain
NLS,: nuclear localization signal
NPC: nuclear pore complex;
PLA: proximity ligation assay
SN: sciatic nerve;
TSA: tyramide signal amplification
WB: western blot;
WES: simple western capillary system

## Supplementary Figures

**Figure S1:**
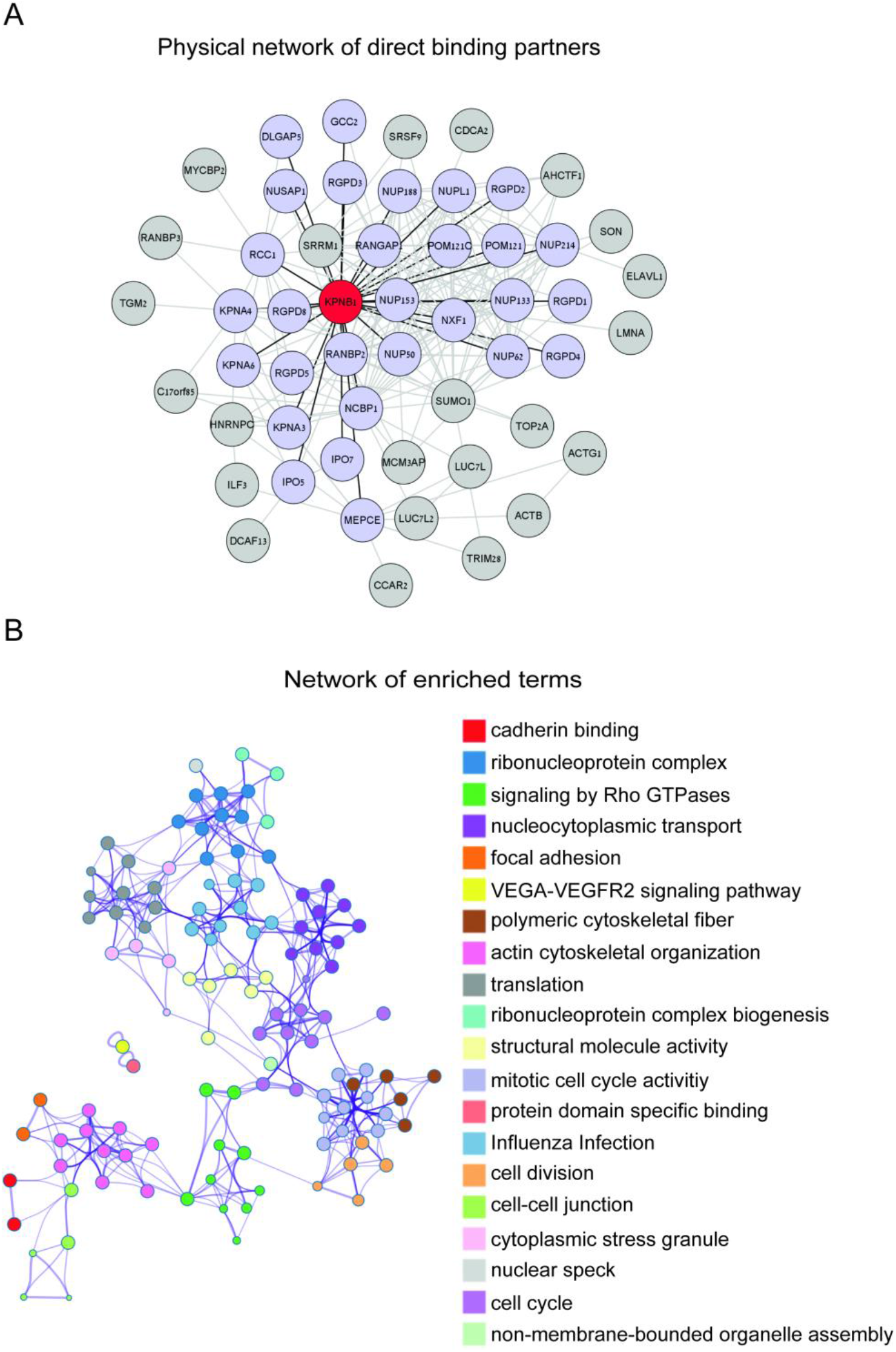
BioID for importin β1 interactors in HeLa cells. (A) Physical network analysis of hits obtained from a combined list of significant candidates from N-KPNB1-BioID and C-KPNB1-BioID constructs showing direct binding partners using STRING with a 0.4 interaction score cut-off value. KPNB1 is highlighted in red. (B) Network of enriched terms: colored by cluster ID, nodes that share the same cluster ID are closer to each other, generated by Metascape.

**Figure S2:**
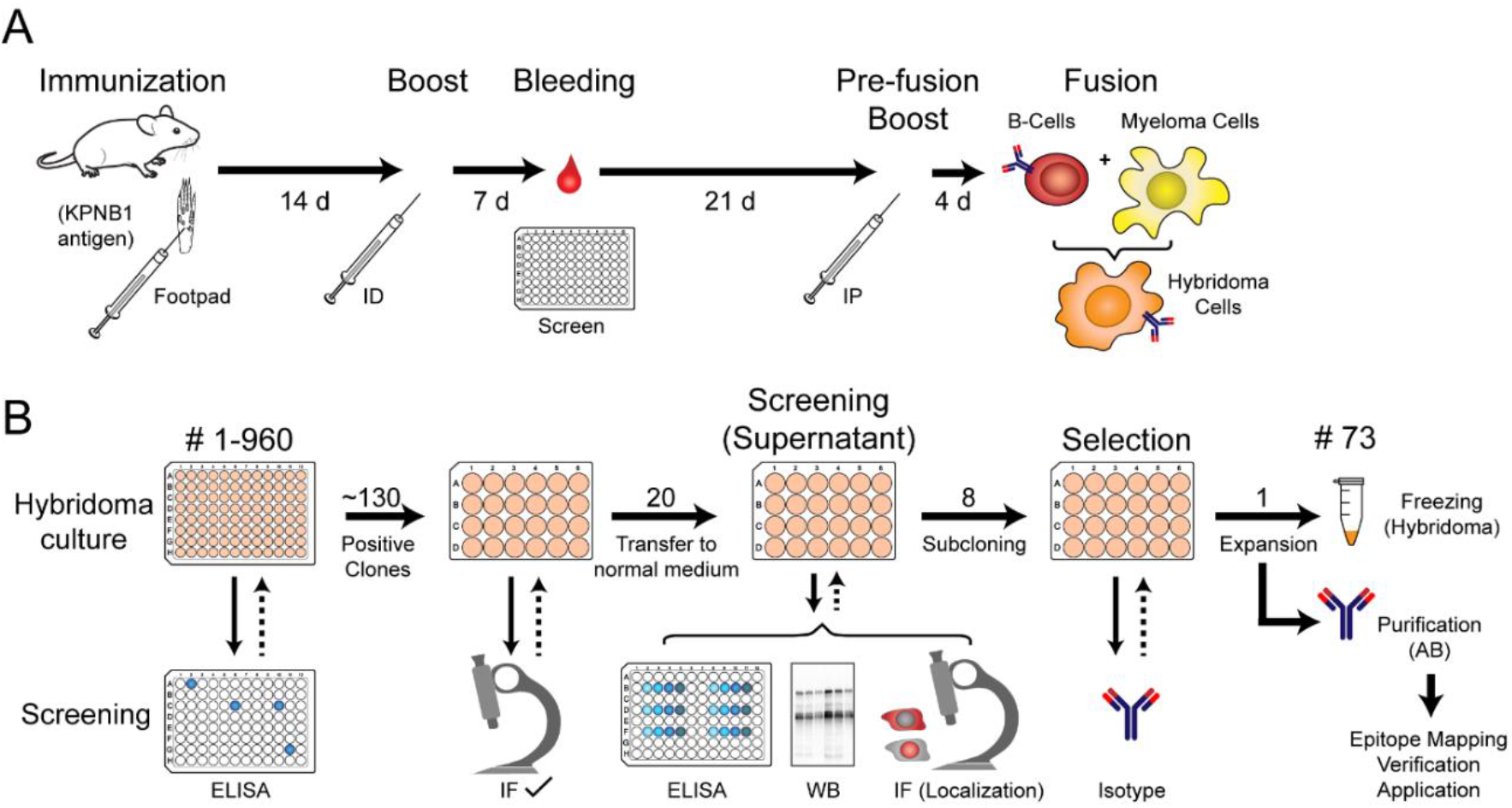
Generation of importin β1 mAbs. (A) Schematic of the immunization timeline. Full-length recombinant mouse importin β1 (KPNB1) protein was used for the immunization via footpad injection. The mouse with the highest immune response (ELISA screen) was used to generate hybridoma cells by fusing spleen cells with myeloma cells. (B) Schematic of the screening procedure and clone selection. Starting with 960 hybridoma cultures, ∼130 clones tested positive for KPNB1 in ELISA and were expanded in 24 well plates. Supernatants were then tested in immunofluorescence (IF) on 3T3 and HeLa cells. The 20 best performing clones underwent further screening and were scored in different applications: linear ELISA, western blot (WB), and IF on DRG neurons. Sub-cloning and isotyping was performed on the top eight clones. Clone #73 was chosen based on total score (suitability for different applications) and its relative specificity for cytoplasmic KPNB1.

**Figure S3:**
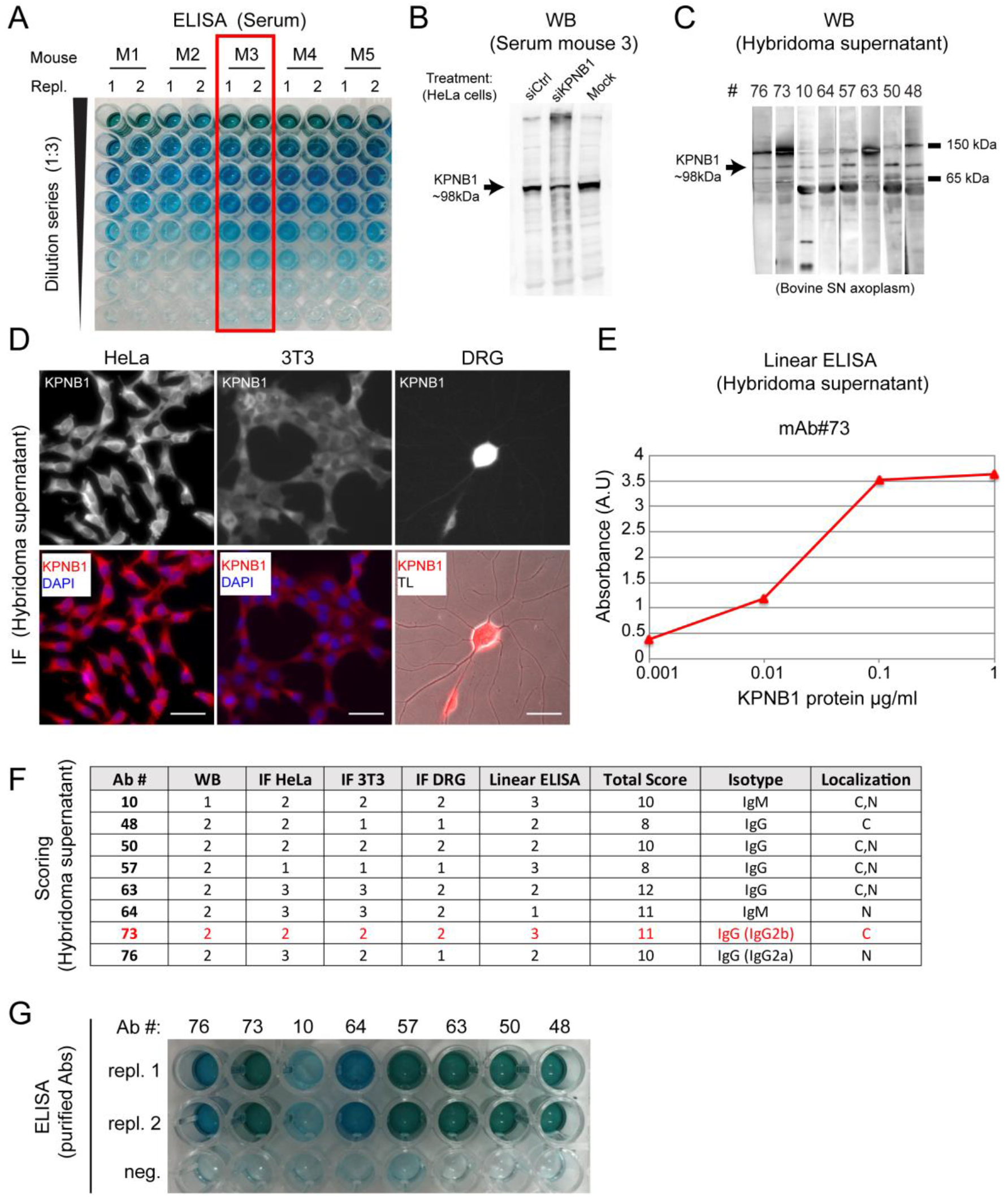
Screening and clone selection. (A) Serum screening of the immunized mice by ELISA for full length KPNB1 reveal Mouse #3 (M3) as the top candidate. (B) KPNB1 Western blot (WB) using serum derived antibodies of M3. (C-E) Screening using cell supernatants shown for selected clones on WB (C), IF, scale bar 20 µm (D), and linear ELISA (E). (F) Summary scores of the best eight clones that were chosen for purification and isotyping. Scores were between 0 (no signal) to 3 (highest signal). Localization of predominantly cytoplasmic or nuclear (C, N) importin β1 is based on IF images. (G) ELISA (full-length mouse KPNB1 with His-tag) using the top eight antibodies after purification. His-tag alone was used as a negative control (neg.).

**Figure S4:**
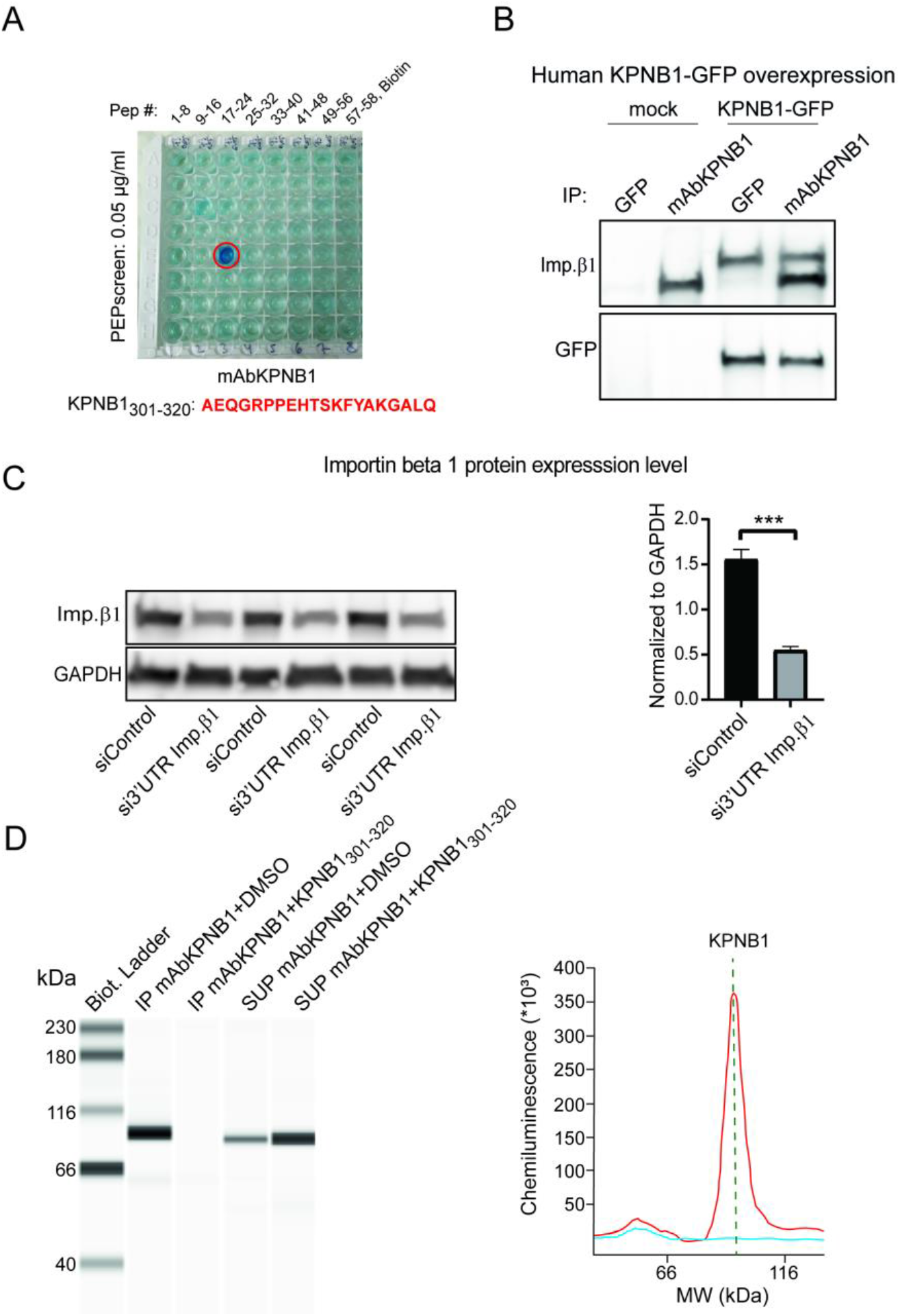
Specificity validation of mAbKPNB1-301-320. (A) Epitope mapping on a custom library comprising 58 biotinylated, overlapping peptides covering the full sequence of KPNB1. Screening was performed in an ELISA format on streptavidin-coated 96-well plates. Peptide P21 comprises residues 301-320 of KPNB1. (B) Immunoprecipitation of ectopically expressed human KPNB1-GFP and endogenous KPNB1 from HEK293T cells. (C) Western blot of siRNA mediated knockdown of endogenous KPNB1 in N2A cells (72 hrs.), KPNB1 levels normalized to GAPDH. Mean ± SEM, Student’s t-test, *** indicates *p* < 0.001. (D) Immunoprecipitation (IP) of KPNB1 from rat brain lysates, using the epitope peptide sequence as a blocking control. Analysis by WES capillary assay is presented as pseudo bands in the left panel and traces in the right panel (red for KPNB1 IP by mAbKPNB1-301-320, light blue in the presence of the blocking peptide).

**Figure S5:**
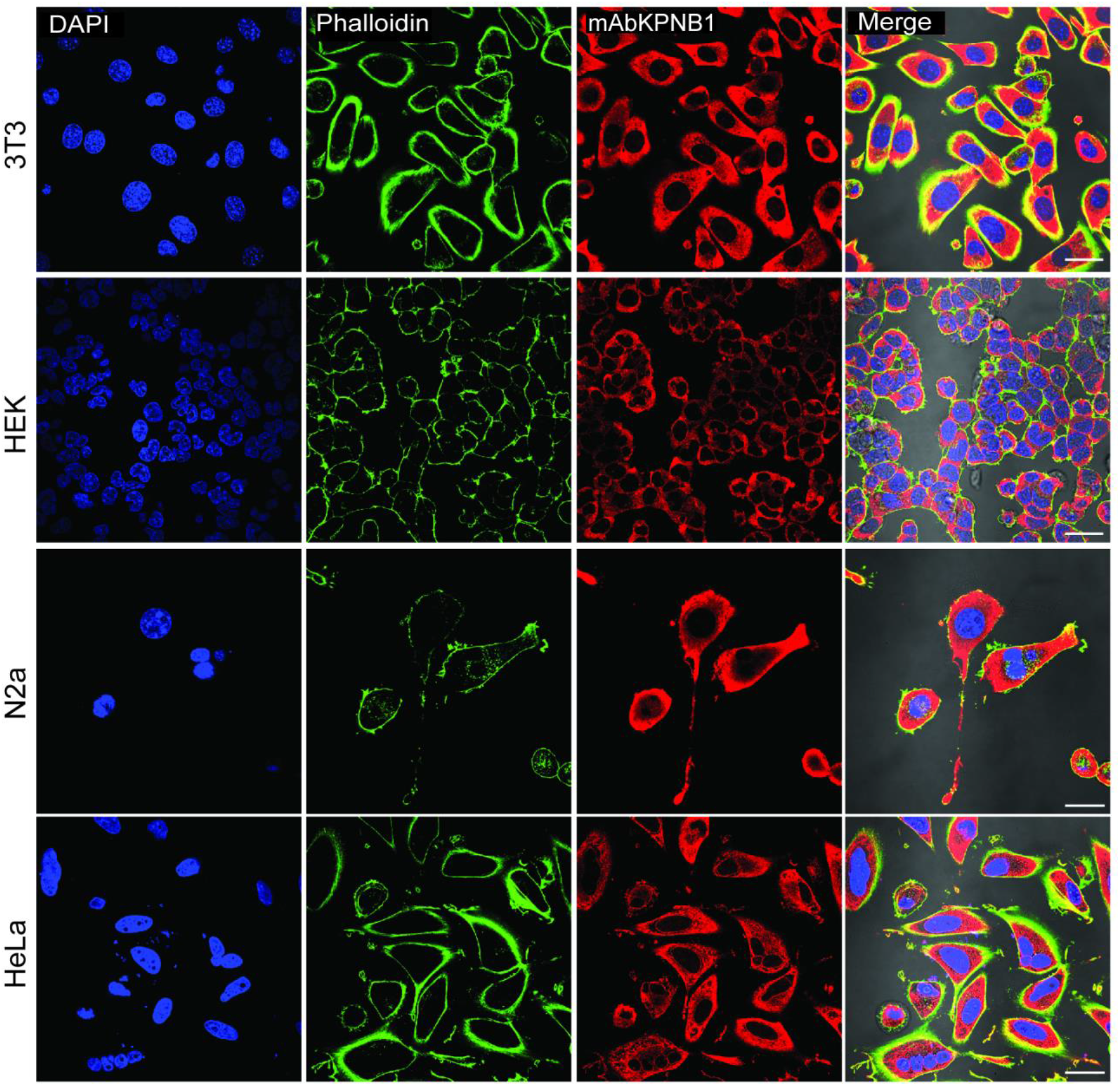
IF in different cell types. Confocal images of 3T3, HEK293T, N2a and HeLa cells stained with DAPI (blue), Phalloidin (green) and 5.4 μg/ml mAbKPNB1-301-320 (red), which shows strong cytoplasmic signal. Merge of all channels shown with transmission light, scale bar 20 μm.

**Figure S6:**
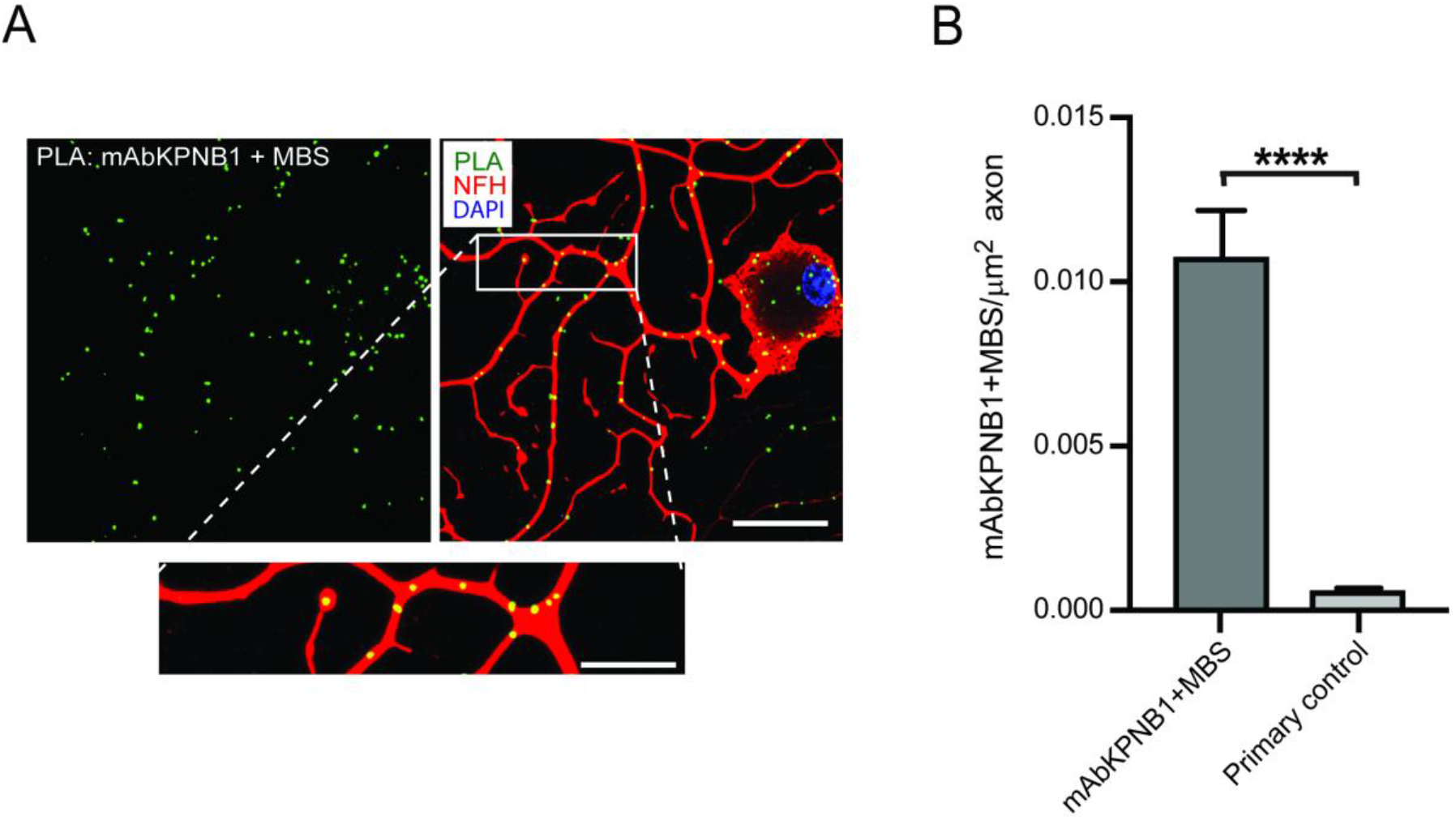
PLA validation of mAbKPNB1-301-320 with commercial KPNB1. **Ab** (A) PLA detection of importin β1 using 2.7µg/ml mAbKPNB1-301-320 and 0.5µg/ml polyclonal anti-KPNB1 (MBS713065) in adult DRG neurons. Scale bars-full image 20 µm; insert = 10 µm. (B) PLA quantification in axons, Mean ± SEM, n ≥ 3, t-test, **** indicates *p* < 0.0001.

**Figure S7:**
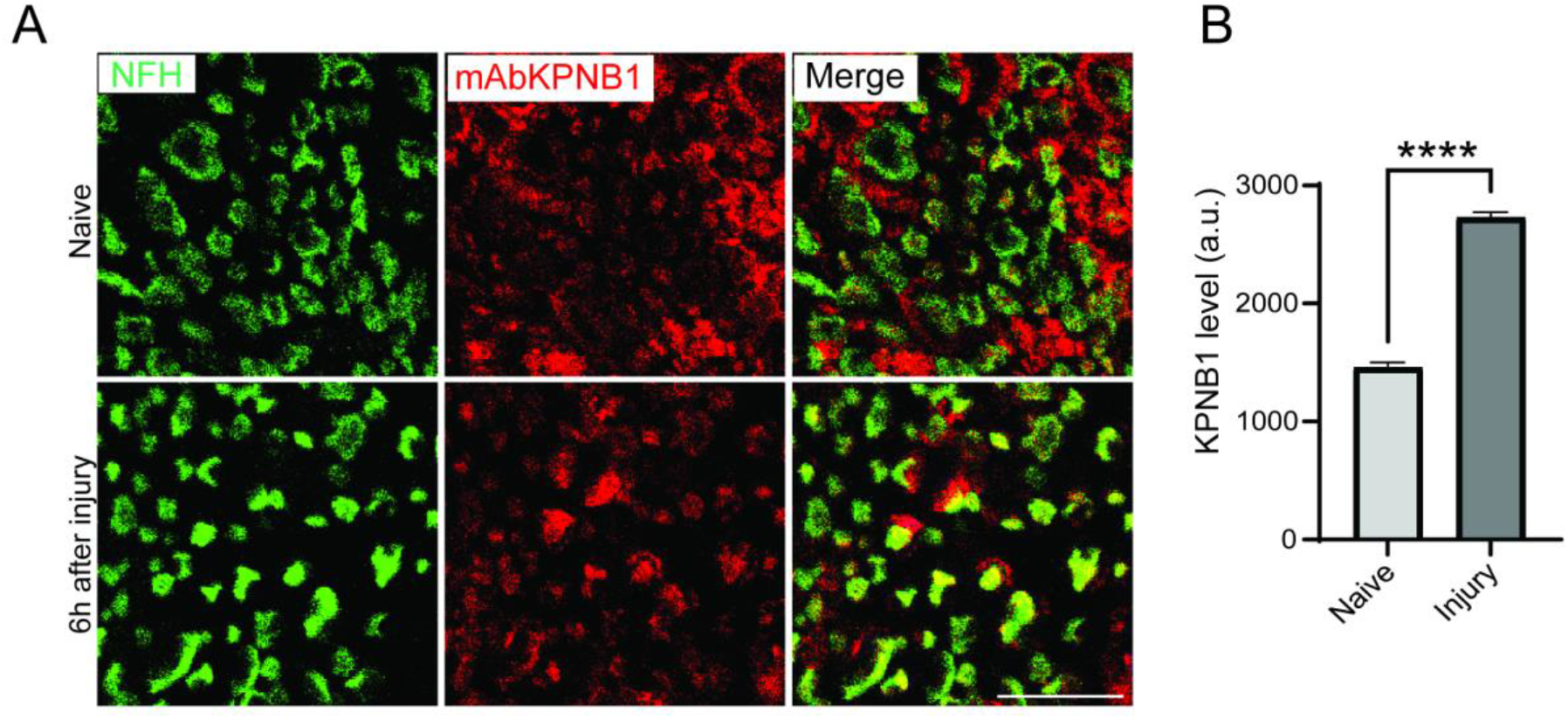
IHC with mAbKPNB1-301-320 on paraffin sections from the sciatic nerve. (A) Paraffin cross sections of the sciatic nerve (SN), showing importin β1 (KPNB1) in red and Neurofilament (NFH) in green. Distance: 5 mm proximal to the site of injury, 6hrs. after SN crush compared to naive. Scale bar 5 μm. (B) Quantification of (A), Mean with SEM, n > 450 axons, unpaired two tailed t-test, **** indicates *p* < 0.0001.

**Figure S8:**
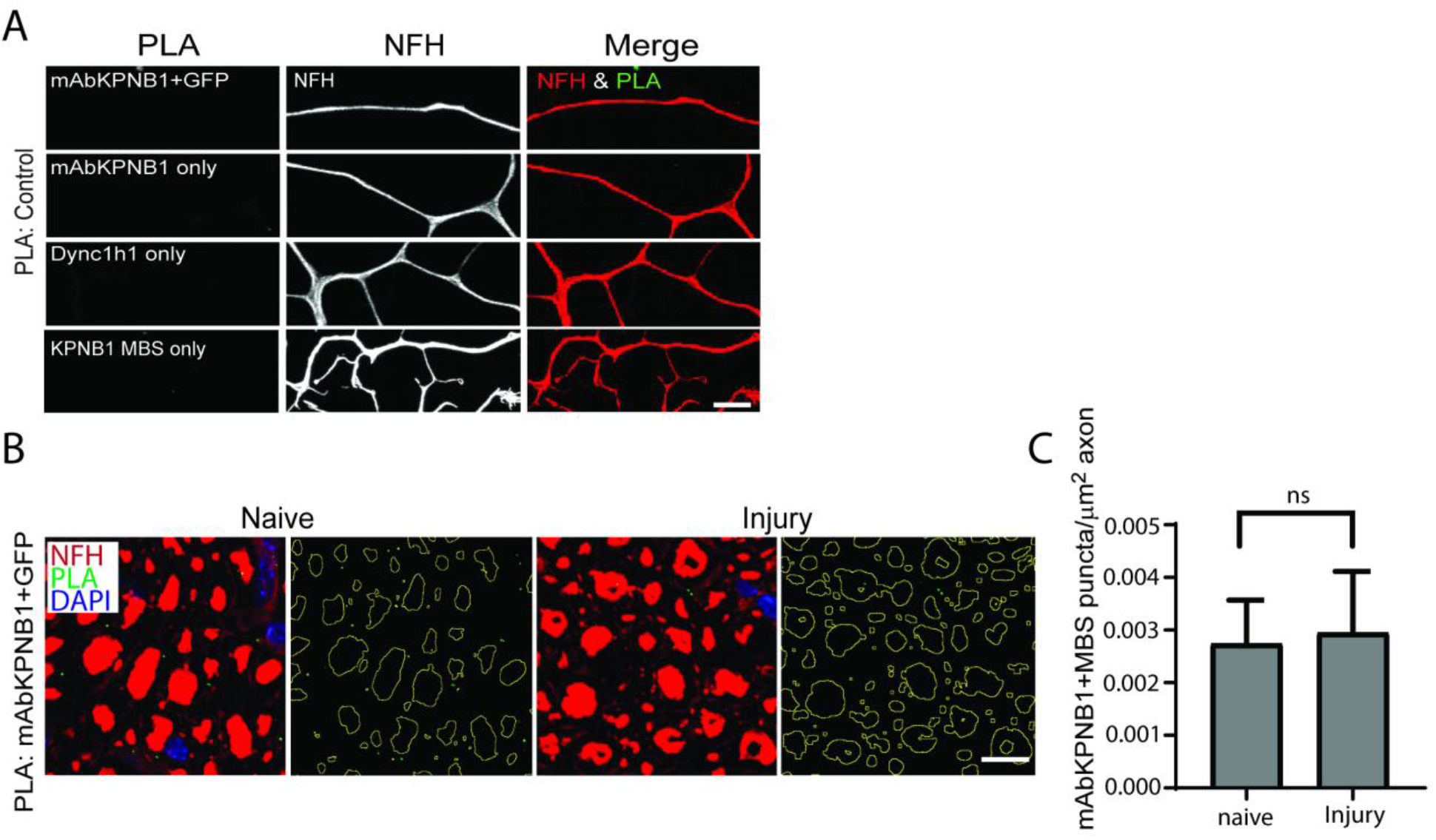
PLA Controls. (A) Confocal images of PLA controls for Figure 4 and Figure S6: mAbKPNB1-301-320 together with rabbit anti-GFP, or single antibody for KPNB1 or Dync1h1. Scale bar 10 µm. (B, C) Confocal images of PLA controls for Figure 5. mAbKPNB1-301-320 together with rabbit anti-GFP (naive N=4, injury N=5). Scale bar 10 µm. Mean ± SEM, unpaired t-test, ns indicates non-significant.

**Figure S9:**
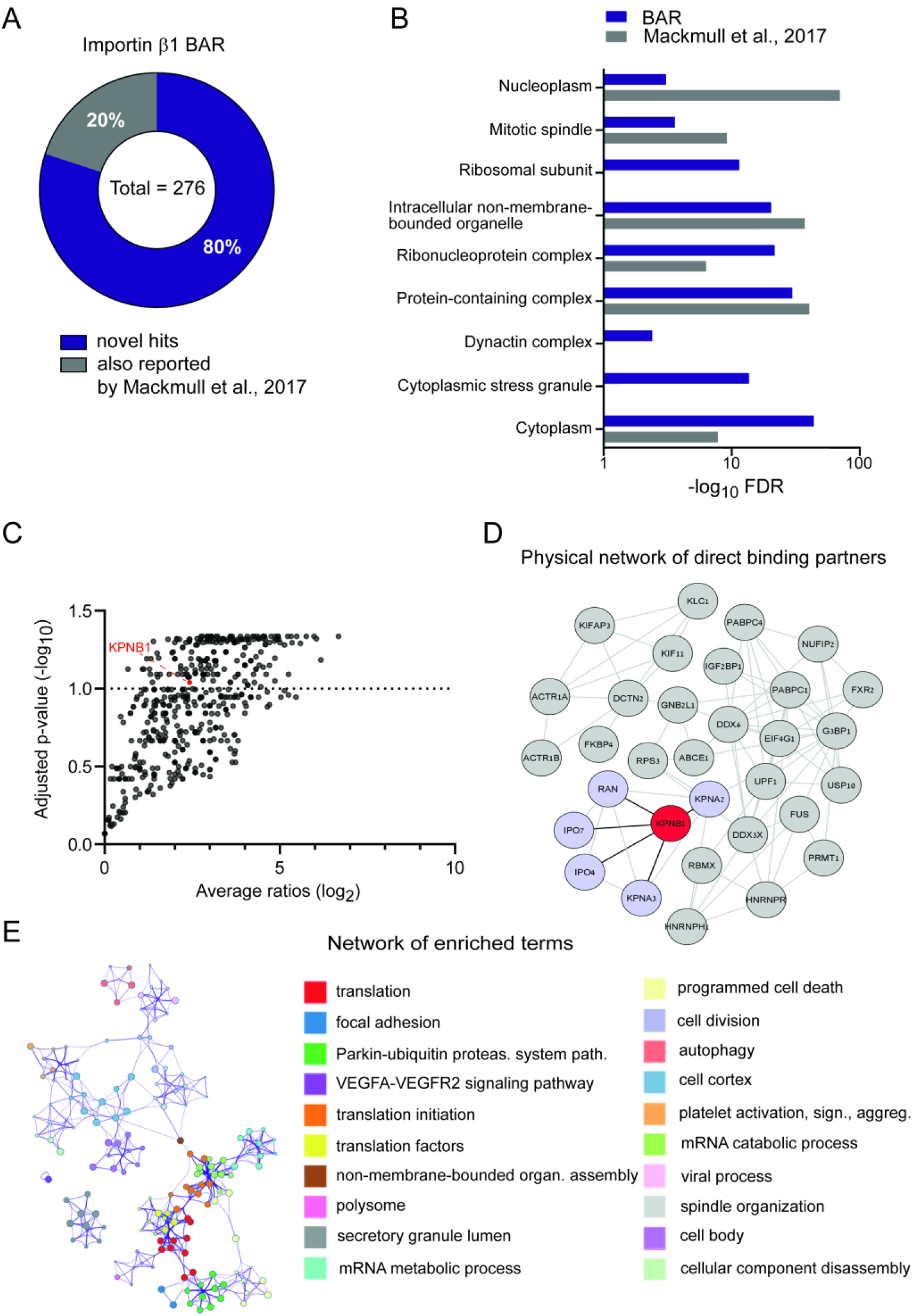
mAbKPNB1-301-320 directed BAR on HEK293T cells. (A) Donut diagram of significant hits from mAbKPNB1-301-320 directed BAR on HEK293T cells, compared to KPNB1 BioID hits from Mackmull et al. 2017. (B) GO analysis using STRING, showing categories for cellular component, p-value cut-off:1%. (C) Volcano plot of proteins identified by mass spectrometry using mAbKPNB1-301-320 directed BAR on HEK293T cells, multiple t-test with a desired false discovery rate of 10%, n=2. (D) Network analysis of hits from (C) showing direct binding partners using STRING database, KPNB1 shown in red, nucleocytoplasmic direct interactors in light purple. (E) Network of enriched terms: colored by cluster ID, nodes that share the same cluster ID are closer to each other, generated by Metascape.

**Figure S10:**
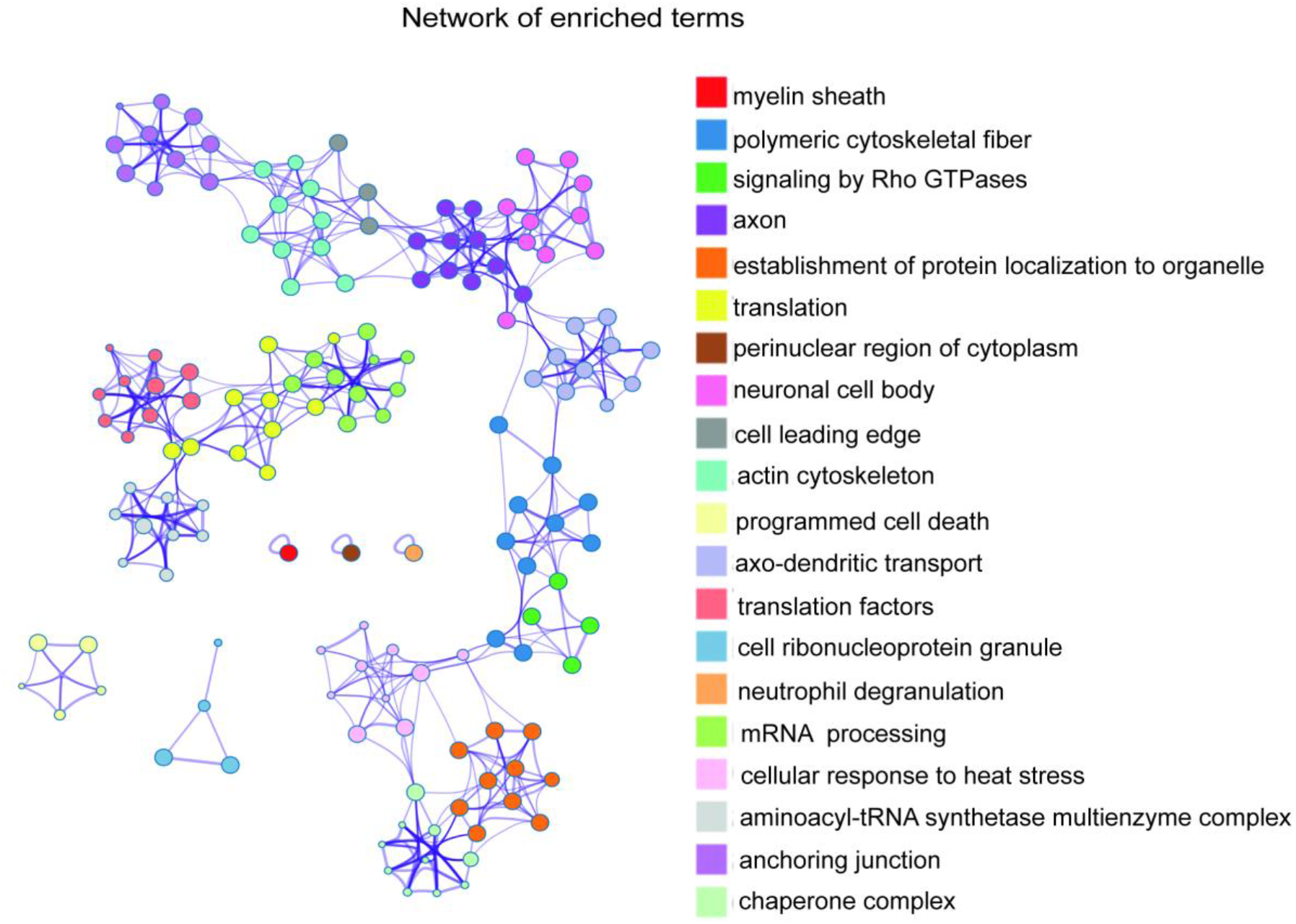
mAbKPNB1-301-320 directed BAR on cultured DRG neurons Extended Figure 6. Enriched terms, colored by cluster ID, nodes that share the same cluster ID are closer to each other, generated by Metascape.

