## Supplementary material for "A new monoclonal antibody enables BAR analysis of subcellular importin β1 interactomes": Methods

### Supplementary Data

#### ***Hybridoma production, antibody screening and purification (extended)***

*Media and Cells*

HBSS: Hank's Balanced Salt Solution

DMEM: Dulbecco's Modified Eagle's Medium, high Glucose 4,5 gr/ml supplemented with 2mM glutamine, 10,000 u penicillin, 10,000 u streptomycin, and 1mM Na pyruvate.

DMEM-HS: DMEM supplemented with 15% heat inactivated (56˚C, 30 min) fetal calf serum.

HAT stocks (100x):
Hypoxanthine (H) $10^{-4}$M ; Stock x 100 = 136 mg/100 ml dist. H2O
Aminopterin (A) 4 x $10^{-7}$M; Stock x 100 = 1.76 mg/100 ml dist. H2O
Thymidine (T) 1.6 x $10^{-5}$M ; Stock x 100 = 38.7 mg/100 ml dist. H2O

HAT-Medium: DMEM-HS supplemented with Hypoxanthine (H), Aminopterin (A) and Thymidine (T) 1:100 from stock

HT-Medium: DMEM-HS supplemented with Hypoxanthine (H) and Thymidine (T)

Polyethylene glycol (PEG): 41% (w/v) in DMEM, sterilized by filtration through 0.45 μm filter or autoclave.

Immune spleen cells: isolated from BALB/c female mice after immunization (Figure S2A). In a petri dish, the spleen was teased apart with sterile forceps in HBSS, and then transferred to a 10 ml tube on ice. The tube was kept in a vertical position for about 3-5 min on ice to let large pieces of tissue (debris) settle down. The supernatant (single cell suspension) was moved to new tube followed by a spin down at 200 g for 10 min. Cells were resuspended in 10 ml DMEM and counted.

Myeloma cells: NS0 cells (cell line derived from the non-secreting murine myeloma, HGPRT(-) variant of MOPC 21 obtained from Dr. C. Milstein). This cell line does not produce any immunoglobulin. It grows in the presence of 20 μg/ml 8-azoguanine (8-AZG) and therefore dies in HAT-medium. NS0 cell line is of H-$2^d$ haplotype and grows in BALB/c mice. Prior to fusion NS0 cells (in logarithmic phase of growth) were counted and resuspend in DMEM at 10 x $10^6$/cells/ml.

*Cell fusion*

Splenic lymphocytes (maximum $10^8$ spleen cells/50ml) and NS0 cells were mixed at a ratio of 5:1 in 50 ml tubes and DMEM was added to a final volume of 40 ml. Cells were centrifuged and the supernatant was completely removed by careful aspiration. Then the tube was flicked gently to loosen the cell pellet, before adding 2ml of prewarmed PEG (drop wise) to the cells. The pellet was resuspended gently for 1 min and incubated for an additional minute at 37˚C. Over the next 15 min, very slowly pre warmed DMEM was added dropwise to dilute the PEG: 5 ml in the first 5 min, then 10 ml in the next 5 min and additional 15-20 ml of DMEM in the last 5 min. After centrifugation and removal of the supernatant, 5 ml of pre warmed DMEM-HS-HAT was added and cells were resuspended very carefully, before adding another 5-10 ml of DMEM-HS-HAT. Finally, viable NS0 or fused cells (bigger than spleen cells) were counted and suspended at a concentration of 1-5 x $10^5$ cells/ml in DMEM-HS-HAT and distributed in 200 µl aliquots into 96 well micro plates and incubated at 37˚C in 8% $CO_2$. In total we obtained 10 x 96-well plates.

Over the next days, cells were monitored for hybrid cell growth and media was replaced after 6 days. The first screening (full protein ELISA) on all 10 plates was done when cells reached ~90% confluency 10 days after plating. Of the 960 colonies, ~130 tested positive in ELISA and were transferred into 24-well plates for expansion and clone selection.

*Screening and clone selection (using hybridoma supernatant)*

During the screening process, 2-3 weeks following fusion, the HAT-medium was replaced by HT-medium and one week later by regular DMEM-HS medium. A schematic of the selection process and hybridoma propagation is shown in Figure S2B. In order to test the positive clones in different applications, the cells were grown up to ~90% confluency and the media (supernatant, containing the AB) was collected and used immediately, or stored at 4°C for up to 2 days.

Since we wanted to analyze subcellular localization and local functions of KPNB1 by Immunofluorescence (IF) we first tested the 130 positive clones for their suitability in IF on cell lines. HeLa and 3T3 cells were grown and fixed in 48-well plates and were subsequently incubated with media from the different hybridoma clones. Antibodies were detected using 488-labeled Fab fragments. Around 20 clones showed robust fluorescent signal and were selected for further characterization with different applications: Linear ELISA (Sensitivity of the AB), Western Blot (Validation of correct molecular weight protein), IF on primary DRG neurons from WT and 3'UTR KO mice (nuclear vs. cytoplasmic/axonal localization). A summary of the results of the screening procedure is shown exemplary for the selected clone #73 (mAbKPNB1-301-320) in Figure S3. The antibodies were scored according to their performance in the different applications - and the following clones were selected for sub-cloning and isotyping: 10, 48, 50, 57, 63, 64, 73, 76, Figure S3F. Scores range from 0 (no signal) to 3 (highest signal).

*Sub-cloning, isotyping, purification and storage*

We started cell cloning under conditions of limiting dilutions (sub-cloning) on our selected clones. Therefore, we made dilutions by serial transfer of 25 µl of cell suspension into 100 µl of medium. Then 100 µl of filler cells (spleen cells, $10^5$ cells/ml) were added. To ensure monoclonality the cloning procedure was repeated for few cycles until all the sub clones detected showed secretion of the specific antibody (via ELISA). An immunoglobulin class/subclass test was performed using a mouse monoclonal antibody isotyping kit, according to the manufacturer's instructions (SBA Clonotyping System-HRP, SouthernBiotech, 5300-05). Clone # 73.25 was selected as the final clone for expansion and subsequent purification. Monoclonal antibodies were purified via Protein A affinity purification, using the AKTA PURE machine. The purified fraction was desalted in PBS, using HiPrep Desalting column and stored at 2-8°C. For long term storage by cryopreservation, ~$10^7$ hybridoma cells were resuspended in 1ml pre-cooled freezing medium (DMEM + 50% Serum + 10% DMSO, sterile by filtration, 4°C). Cells were frozen slowly, kept for 48h at -80°C and then moved to liquid nitrogen.

All procedures were based on protocols adapted from Eshhar, Z. (1)

#### *Immunohistochemistry for paraffin sections*

Sciatic nerves from naive and injured mice (6h) were fixed in 4 % PFA overnight at 4 °C, dehydrated by increasing concentrations of ethanol and subsequently embedded in paraffin blocks. For analyzing importin-β1 in naive and injured nerves, 5 µm cross sections were taken at different locations proximal to the crush site. The slides then underwent deparaffinization with xylene and ethanol, followed by antigen retrieval in Tris-EDTA buffer (10 mM Tris Base, 1 mM Ethylenediaminetetraacetic Acid (EDTA), 0.05 % Tween20, pH 9.0) using a pressure cooker at 125 °C for 1 min with subsequent cooling to RT for 1 h, and 2 washes in PBS. Blocking was performed in 20 % horse serum, 0.2 % Triton in PBS for 1 h at RT. Primary antibodies (mAbKPNB1 2µg/ml, NFH 1:1000) were incubated in Ab-solution (2 % horse serum, 0.2 % Triton in PBS) over night at RT. Secondary Abs (Donkey, 1:500, Jackson ImmunoResearch) were applied in Ab-solution for 2 h at RT. Slides were mounted in Fluoromount (F4680, Sigma Aldrich) and confocal images were taken using an Olympus FV1000 Confocal laser-scanning microscope at 60x magnification with oil-immersion objective (Olympus UPLSAPO, NA 1.35).

##### *Cloning of BioID-constructs*

All cloning steps were carried out using the restriction-free (RF) cloning protocol as previously described. (2, 3). The KPNB1 CDS was amplified from pEGFP-N1 human full length KPNB1 (Addgene #106941) and mVenus CDS was amplified from pTrix-mVenus-PA-Rac1, Addgene #22007). The multiple cloning site of pcDNA3.1 mycBioID (Addgene #35700) was removed using primers MCSout-fwd and MCSout-rev to generate Myc-BirA*.

<u>Myc-BirA*-ImpB1-mVenus construct:</u> The KPNB1 CDS was amplified using primers BirA*-ImpB1-fwd and BirA*-ImpB1-rev and inserted into Myc-BirA* to generate plasmid Myc-BirA*-ImpB1. mVenus CDS was amplified using primers ImpB1-mVenus-fwd and ImpB1-mVenus-rev and inserted into Myc-BirA*-ImpB1 resulting in plasmid Myc-BirA*-ImpB1-mVenus.

<u>Myc-mVenus-ImpB1-BirA* construct:</u> The KPNB1 CDS was amplified using primers ImpB1-BirA*fwd and ImpB1-BirA*rev and inserted into Myc-BirA* to generate plasmid Myc-ImpB1-BirA*. mVenus CDS was amplified using primers mVenus-ImpB1-fwd and mVenus-ImpB1-rev and inserted into plasmid Myc-ImpB1-BirA* resulting in plasmid Myc-mVenusB-ImpB1-BirA*.

| plasmid/construct | primer | sequence 5'-3' |
| --- | --- | --- |
| Myc-BirA* | MCSout-fwd | TCCCTGAGAAGCGCAGAGAAGCTCGAGCGGTGATCAGCCTCGACTGTGCCTTCTAGTTGC |
| | MCSout-rev | GCAACTAGAAGGCACAGTCGAGGCTGATCACCGCTCGAGCTTCTCTGCGCTTCTCAGGGA |
| Myc-BirA*-ImpB1-mVenus | BirA*-ImpB1-fwd | GGAGAAATCTCCCTGAGAAGCGCAGAGAAGGGATCCGAGCTGATCACCATTCTCGA |
| | BirA*-ImpB1-rev | AGGCACAGTCGAGGCTGATCACCGCTCGAGTCAAGCTTGGTTCTTCAGTTTCCTC |
| | ImpB1-mVenus-fwd | CAAAAGAACTGAGGAAACTGAAGAACCAAGCTGGATCTGTGAGCAAGGGC |
| | ImpB1-mVenus-rev | CACAGTCGAGGCTGATCACCGCTCGAGTCACTTGTACAGCTCGTCCATGC |
| Myc-mVenus-ImpB1-BirA* | ImpB1-BirA*-fwd | GAACAAAAACTCATCTCAGAAGAGGATCTCGACGAGCTGATCACCATTCTCGA |
| | ImpB1-BirA*-rev | GATCAGCTTCAGGGGCACGGTGTTGTCCTTGGATCCAGCTTGGTTCTTCAGTTTCCTC |
| | mVenus-ImpB1-fwd | GAACAAAAACTCATCTCAGAAGAGGATCTCGACGTGAGCAAGGGCGAGGAG |
| | mVenus-ImpB1-rev | CACGGTCTTCTCGAGAATGGTGATCAGCTCAGAACCCTTGTACAGCTCGTC |

#### ***References***

1. Eshhar, Z. (1985) Monoclonal Antibody Strategy and Techniques. In: Springer, T. A., ed. *Hybridoma Technology in the Biosciences and Medicine*, pp. 3-41, Springer US, Boston, MA

2. Peleg, Y., and Unger, T. (2014) Application of the Restriction-Free (RF) cloning for multicomponents assembly. *Methods Mol Biol* 1116, 73-87

3. Unger, T., Jacobovitch, Y., Dantes, A., Bernheim, R., and Peleg, Y. (2010) Applications of the Restriction Free (RF) cloning procedure for molecular manipulations and protein expression. *J Struct Biol* 172, 34-44
